# Mesoscale functional architecture in medial posterior parietal cortex

**DOI:** 10.1101/2023.08.27.555017

**Authors:** Riichiro Hira, Leah B. Townsend, Ikuko T. Smith, Che-Hang Yu, Jeffrey N. Stirman, Yiyi Yu, Spencer LaVere Smith

## Abstract

The posterior parietal cortex (**PPC**) in mice has various functions including multisensory integration^1–3^, vision-guided behaviors^4–6^, working memory^7–13^, and posture control^14,15^. However, an integrated understanding of these functions and their cortical localizations in and around the PPC and higher visual areas (**HVAs**) has not been completely elucidated. Here we simultaneously imaged the activity of thousands of neurons within a 3 x 3 mm^2^ field-of-view, including eight cortical areas around the PPC, during behavior with a two-photon mesoscope^16^. Mice performed both a vision-guided task and a choice history-dependent task, and the imaging results revealed distinct, localized, behavior-related functions of two medial PPC areas. Neurons in the anteromedial (**AM**) HVA responded to both vision and choice information, and thus AM is a locus of association between these channels. By contrast, the anterior (**A**) HVA stores choice history with sequential dynamics and represents posture. Mesoscale correlation analysis on the intertrial variability of neuronal activity demonstrated that neurons in AM exhibited diverse, area-dependent interactions, while neurons in area A shared fluctuations with the primary somatosensory area. Pairwise interareal interactions among neurons were precisely predicted by the anatomical input correlations, with the exception of some global interactions. Thus, the medial PPC has two distinct modules, areas AM and A, with each having distinctive modes of cortical communication. These medial PPC modules can serve separate higher-order functions: area AM for multisensory and cognitive integration with locally processed signals and area A for transmission of information including posture, movement, and working memory.

## INTRODUCTION

The posterior parietal cortex (PPC) is an association area located between the visual and somatosensory cortices and is widely shared by mammals. The PPC has distinct regions and each one of these has unique functions and corticocortical projection patterns in primates ^17–19^. In mice, the PPC overlaps with medial and anterior HVAs ^20^, has characteristic cytoarchitectures ^21–24^ and unique corticocortical recurrent projection patterns ^4,7,22^, which are similar to those of rats ^25,26^. Despite its relatively small area compared to that of primates, mouse PPC has various functions like that of primates including navigation ^7,8,10–12,27,28^, multisensory integration ^1–3^, memory-guided decision making ^6–9,29^, visually ^4,5^ and auditory ^9,30^ guided decision making. In rats, PPC is also important in attention ^31,32^, sensory history ^13^, and posture coordination ^14,15^. Thus, neurons in rodent PPC can represent various sensorimotor and integrative functions. However, prior studies typically focused on a single cortical location and often a single function, and thus it is unclear whether neurons with particular functions are distributed in overlapping populations or localized.

The function of an individual cortical area is influenced by the information that area receives from other brain regions. Connectivity among cortical areas is highly specific in mice ^33^, and includes frequent cases of neurons sending bifurcated axons to multiple target regions ^34^. This axonal divergence indicates that to understand association inter-areal interactions, it is insufficient to consider only pairwise connections. Instead, such analysis should consider the shared connectivity patterns among three or more areas. When weak connections are included, the mouse cerebrum is estimated to have a 97% connectivity density ^35^. In addition, cortical activity is a complex interplay of diverse information due to interactions with the thalamus, basal ganglia, and cerebellum. Moreover, the mode of these interactions can vary depending on behavior ^36,37^. Thus, to understand the nature of the PPC, it is necessary to examine not only the pairwise interactions between cortical areas, but also investigate multiple interrelated areas in an integrated manner under a variety of environmental constraints.

Here, we examine the functional architecture of the mouse PPC and surrounding cortical areas using calcium imaging with a large field-of-view two-photon mesoscope ^16^ during head-fixed behavior in multiple tasks. We mapped vision, choice, choice history, and posture-related neurons within and surrounding the PPC. We found clear functional localization between two areas in medial PPC: HVAs A and AM. Furthermore, the mesoscale correlation structure of trial-to-trial variability indicated that A and AM had distinct relationships with surrounding cortical areas. The local interareal interaction aligned with the anatomical structure, but the global interaction did not, and thus the functional data revealed a dynamic aspect of the circuit organization. Our findings indicate that various modes of interareal communication constitute diverse functions of PPC.

## RESULTS

### Demarcation of PPC as A, AM, and RL

To functionally determine the precise anatomical location of PPC, we first identified retinotopic and somatotopic organizations with intrinsic signal optical imaging (ISOI) in anesthetized mice ^38–42^. Retinotopic mapping revealed locations of each HVA including AM (**Figure 1A**), which has been considered to be the core of mice PPC in recent studies ^4,27^. To map the boundary of the primary somatosensory (**S1**) area anterior to the HVAs, we used tactile vibration of the body parts including the trunk, as trunk S1 is known to be located at the most posterior portion of the S1 ^33^. As expected, the trunk S1 (**S1t**) was located at a posterior region within S1 (**Figure 1B; Figure S1A,B**). The distance from the center of the AM to the center of the S1t was ∼900 μm (894 ± 18.3 μm (mean ± s.d.), n = 8 mice). Assuming that the diameter of AM and S1t are 600 μm and 400 μm, respectively, we could estimate that there would be a space 400 μm wide along the rostrocaudal axis between these two areas. Here, we defined this space as area A (**Figure 1C**), because it roughly overlaps with the HVA identified as area A in a landmark anatomical study of mouse HVAs ^20^. Our definition of area A likely includes a larger area than this original definition.

**Figure 1.**
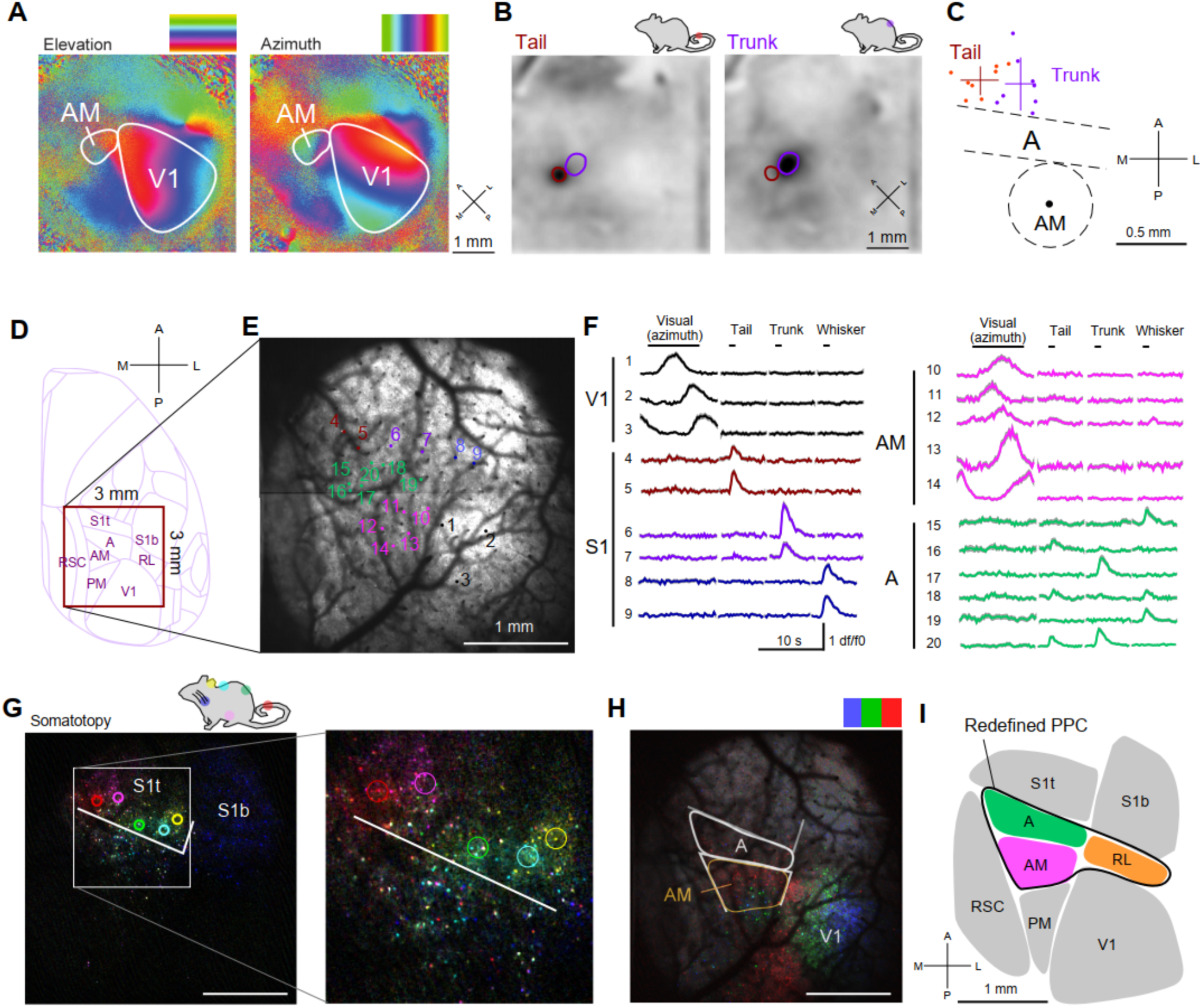
**Definition of cortical area A and redefinition of PPC based on visual and somatosensory responses. A.** Elevation *(left)* and azimuth (*right*) maps showing retinotopy that form the borders of V1 (primary visual cortex) and AM. The cortical surface was color-coded according to the location of the bar on the monitor at the time of the evoked visual response. **B.** Tactile stimuli of a tail (left, red) and a trunk (*right*, green) resulted in clear spots in S1. **C.** The center positions of S1 tail and S1 trunk regions relative to the center position of AM from 8 mice. The gap space between the S1 trunk region and AM was defined as A. **D.** A location of the cranial window and FOV of two-photon imaging was overlaid on the Allen CCF version 3 ^33^. **E.** The average image of calcium imaging and ROI positions for panel **F**. **F**. Fluorescence traces (mean and s.e.m.) of ROI indicated in panel **E** when the mouse received visual stimulation (azimuth) and tactile stimuli (tail, trunk, and whisker). **G,H.** Single neuron resolution mapping of somatotopy (**G**) and retinotopy (**H**) revealed that A had neither retinotopy nor somatotopy, but had neurons activated by tactile stimuli and a small number of neurons activated by visual stimulation. On the other hand, AM had retinotopy and a small number of neurons activated by tactile stimuli. A was defined after the S1 and AM were determined by somatotopic and retinotopic organizations, respectively. Scale bar: 1mm. **I**. Redefined PPC in this study (A, AM and RL). Abbreviations: RL, rostrolateral HVA; PM, posteromedial HVA; RSC, retrosplenial cortex.

Often, PPC is determined using stereotaxic coordinates, and we considered this approach. However, systematic mapping of visual areas relative to the lambda and midline showed that each HVA location based on the stereotaxic coordinates was unreliable. The mean distance from the center position of each HVA in each mouse to the average center position across mice was 244 μm, (range: 198 – 316 μm, n = 12 mice) (**Figure S1C,D**) when the positions were measured relative to the lambda landmark. We confirmed that the variability in locations was not explained by the age or sex **(Figure S1E**). Instead, the variation of HVAs substantially decreased after the registration of the areas by translation based on the location of V1 and rostrolateral area (RL). After this registration, the mean distance from the center position of each HVA to the average center position across mice was ∼91.5 μm (range: 38.6 - 141 μm), n = 12 mice): a reduction in variability by more than half **(Figure S1F-I)**. This improvement indicates that relative locations of HVAs are similar across mice ^43^, but HVA locations relative to skull landmarks are more than twice as variable. Thus, localizing these small HVAs and components of PPC by stereotaxic coordinates can introduce variability and decrease the fidelity. We concluded that functional imaging of retinotopic and somatotopic organization must be obtained in each individual mouse to precisely and rigorously map HVAs and PPC, and to ascribe functions to neurons therein.

ISOI was supplemented with two-photon calcium imaging to observe the activity of neurons within and surrounding PPC including area A (**Figure 1D-F**). In awake mice, somatosensory stimuli including tail, trunk, neck, and whiskers resulted in responses in PPC areas. In S1, somatosensory stimulation evoked localized neuronal responses that exhibited somatotopic organization. In contrast, area A neurons activated by somatosensory stimuli did not exhibit clear somatotopy (**Figure 1G**). Likewise, visual responses of neurons in area A did not exhibit clear retinotopy (**Figure 1H**), in contrast to other HVAs ^20^. Some area A neurons showed activity by stimulation of multiple body parts such as trunk and tail, or trunk and whiskers, which is similar to area 5 of macaque ^44^ (**Figure 1F**). Thus, area A integrates multiple somatosensory inputs, potentially contributing to body coordination (e.g., posture) ^14,44^. Some neurons in AM were activated by tactile stimuli, and again this activity lacked somatotopy. Based on the multisensory responsiveness of areas A, AM, and RL ^2,45,46^, we demarcate the PPC as a three-module structure, consisting of AM, RL, and A (as defined above) (**Figure 1I**).

### Visually guided task

After precisely mapping PPC and adjacent cortical areas, we examined layer 2/3 neuronal activity during behavior with single neuron precision across multiple cortical areas simultaneously with large field-of-view, two-photon calcium imaging (**Movie S1**). We used transgenic mice that express a genetically encoded calcium sensor, GCaMP6s, in cortical neurons ^47^. The field-of-view (**Figure S2**) typically covered PPC and surrounding areas including HVAs, V1, S1, and a portion of retrosplenial cortex (RSC).

PPC has been reported to have functions including visually guided decision-making and working memory. Do these functions differ among A, AM and RL? To address this question, we trained the mice under head-fixed conditions to perform vision-guided and history-guided two-alternative forced-choice tasks (**Figure S3A-D**) and imaged the activity of neurons for each task. First, we investigated the distribution of task-related neurons during a visually-guided task (“**vision task**”; **Figure 2A-D**). In this task, mice had to lick one of two waterspouts (left or right) based on the visual stimuli, naturalistic video (**Movie S2**) or black screen, presented to the left visual field of the mice (imaging in the right hemisphere) (**Figure S3A,B**). This simple task was designed to clearly map sparse responses of neurons to natural video ^48^, and to map the choice-related activity guided by the visual stimuli at the same time to see segregation or overlapping of the vision and choice-related neurons around PPC.

**Figure 2.**
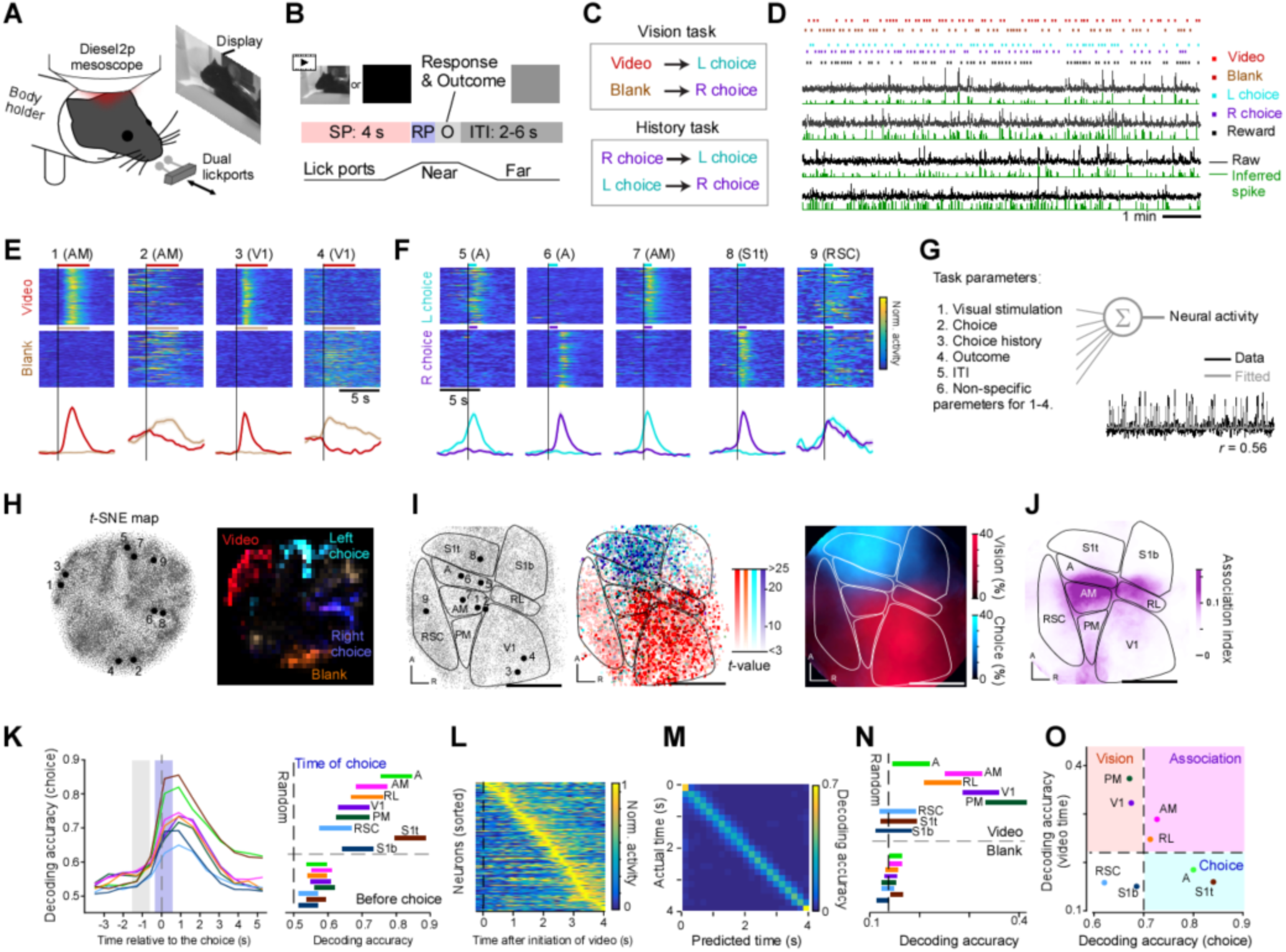
**Vision and choice information are merged in AM. A.** Schematic illustration of a vision-guided choice task. **B.** The lick-ports were approached after a 4-s sampling period (SP). The mouse was given a reward (water) or a punishment (air-puff) based on the licking direction during response period (RP). ITI, Inter-trial-interval. **C.** Rules of vision and history task. **D.** Four example raw traces (black) and inferred spikes (green) during the task. **E.** Representative vision neurons (ROI 1-4 in **I**). The red bars indicate sampling periods during video presentation, and the brown bars indicate sampling periods without video stimulation. Vertical black lines mark the onset of the sampling period. **F.** Representative choice neuron (ROI 5-8 in **I**) and a non-selective neuron (ROI 9). Light blue lines indicate the response periods in trials with left choices, and purple lines indicate the response periods in trials with right choices. Vertical black lines mark the onset of the response period. **G.** Schematic of our encoding model. The bottom right panel shows an example of single-neuron activity with an overlay of the fitting obtained by the encoding model. **H.** *t*-SNE result of single neuron activities (left). Task-related neurons were mapped onto the *t*-SNE (right). **I**. All active neurons in the vision task (left) and the *t*-values of video (red), blank (orange), left choice (cyan), and right choice (purple) are overlaid (middle). The ratio of vision and choice neurons is also shown (right). **J.** Map of Association index (harmonic mean of vision and choice neurons.) **K**. Decoding accuracy of choice by activity of 50 neurons from eight areas before and after the choice were plotted (left). The decoding accuracies 1s before (right bottom, gray shade; p=0.13; one-way ANOVA) and just after (right top, blue shade; p=3.6 × 10^--9^; one-way ANOVA) the choice was shown. **L.** The activity of vision neurons was averaged and aligned by the order. **M.** The time after the onset of the video was precisely decoded with a population of vision neurons in panel **L. N.** Decoding accuracy of time during video presentation (top; p=1.1 × 10^-15^; one-way ANOVA) and black screen presentation (bottom; p=0.013; one-way ANOVA) were plotted. **O.** Decoding accuracies of time in video presentation and choice direction indicate that AM would be the best position for associating these two signals. The background color and dashed lines are provided as visual aids for illustrative purposes. Error bars, mean ± s.e.m. in **E** and **F**, 95% confidence interval in **K** and **N**. Scale bar: 1mm.

Large field-of-view calcium imaging during the task enabled us to simultaneously observe ∼10^4^ neurons (n = 5 mice, total of 10 sessions, 47162 neurons). Neurons were robustly activated by specific frames in the naturalistic video, by the black screen, or by left or right choice (**Figure 2E,F; Movie S3**). Notably, some PPC neurons showed complex dynamics rather than simply encoding specific visual or choice information, indicating integrative roles. To quantitatively identify and map the functional properties of each neuron, we constructed a simple linear regression model (**“encoding model”, Figure 2G**). In the encoding model, the activity of each neuron was fitted by a weighted sum of external control parameters, such as video frames, and behavioral parameters, such as choice and reward direction. Because the visual stimulus changes continuously over time, sliding time windows were placed during the visual stimulus period. The linear model was sufficient to estimate the correlation with neural activity since there is little correlation between task parameters such as visual stimuli, choice, reward, and inter-trial-intervals (**materials and methods**). The statistical significance (p<0.001) of each parameter corresponding to visual stimulus and chosen direction was used to identify the vision and choice-related neurons, respectively. To examine whether the neuron groups labeled by this model broadly capture the diversity of neuronal activity, we performed unsupervised clustering of neuronal activity using t-SNE. The functional labels revealed by this encoding model were consistent with the *t*-SNE clusters, indicating the validity of the encoding model (**Figure 2H; Figure S4B; materials and methods**). The neurons which were selective to video or black screen (**“vision neurons”**) were found in V1 and HVAs including AM, while the neurons which were selective to the chosen direction (**“choice neurons”**) were distributed in S1t and A (**Figure 2I**). Interestingly, the neurons that activated during the choice period irrespective of the choice directions were distributed broadly (**Figure S4A**). The vision and choice neurons were distributed mostly exclusively, but some overlap was found around AM as revealed by mapping of the harmonic mean of the percentage of vision and choice neurons (**“association index”**) (**Figure 2J; Figure S4A; Movie S3**). This indicates that the AM would be suitable for integrating vision and choice information rather than the A, RL or the other areas around PPC.

The PPC neurons had been thought to be important for decision making, but recently it has also been suggested that the PPC neurons store the history information rather than deciding the upcoming choice. To analyze whether the neurons around PPC represent the upcoming choice or the past choice in our visually guided task, we decoded the choice direction with the activity of 50 neurons before and after the choice (**Figure 2K**). The decoding accuracies of A and S1t were higher than the other areas at the choice timing and they stored the choice information even 5 s after the choice. On the other hand, the decoding accuracy of the choice 1 s before the choice was very low in any areas recorded. Thus, neurons in areas A and S1t reflect the choice direction immediately after the choice and store it over 5 s. In addition, vision neurons preferred a specific time of the video frame resulting in the vision neurons sequentially firing during the sampling period (**Figure 2L**). Decoding analysis showed that the vision neurons precisely possessed the temporal information of the video frame (**Figure 2M**). Such precise decoding of time during the sampling period was not observed in black screen trials (**Figure 2N**). From the two decoding performances, we divided the imaged areas into four groups (**Figure 2O**). Note that area AM is the most suitable area for the association of these two parameters, as it possesses a substantial amount of decodable information about both parameters.

### Choice history guided task

Next, we asked whether and where choice history information can be stored in PPC and surrounding areas. We designed a choice history-dependent task (“**history task**”) with the same set up as the vision task (**Figure 2A-C**). In this task, the mice had to choose the opposite side of the previous choice. Mice had to store the direction of the choice for 6-10 seconds from the previous choice to the next choice including an inter-trial-interval (**ITI**) and a sampling period. Since the visual stimuli were randomly presented during the sampling period, the mice had to ignore the visual stimuli. This task was designed to map the history neurons in contrast to the choice neurons and the vision neurons around PPC.

Large FOV calcium imaging during the history task (n=4 mice, 10 sessions, 27059 neurons) revealed that neurons in A and S1t maintained their activity during the ITI in a past-choice-specific manner, indicating that these neurons stored the choice history for making the next choice (**Figure 3A; Figure S3G**). We used the same encoding model as in the vision task to identify the neurons representing the history of the previous choice (**“history neurons”**) as well as choice and vision neurons. Again, the functional labels revealed by this encoding model were consistent with the *t*-SNE clusters, indicating the validity of the encoding model (**Figure 3B; Figure S4C,D**). We found that neurons in area A had a significantly larger ratio of history neurons than all the other areas including S1t that had more choice neurons, and that neurons in AM exhibited large association index (**Figure 3C,D**). We also found more history neurons in the history task in A compared to those in the vision task (**Figure 3E**). Thus, A exhibited a task-specific functional change, preferentially representing a parameter when it is relevant, despite being available in both tasks.

**Figure 3.**
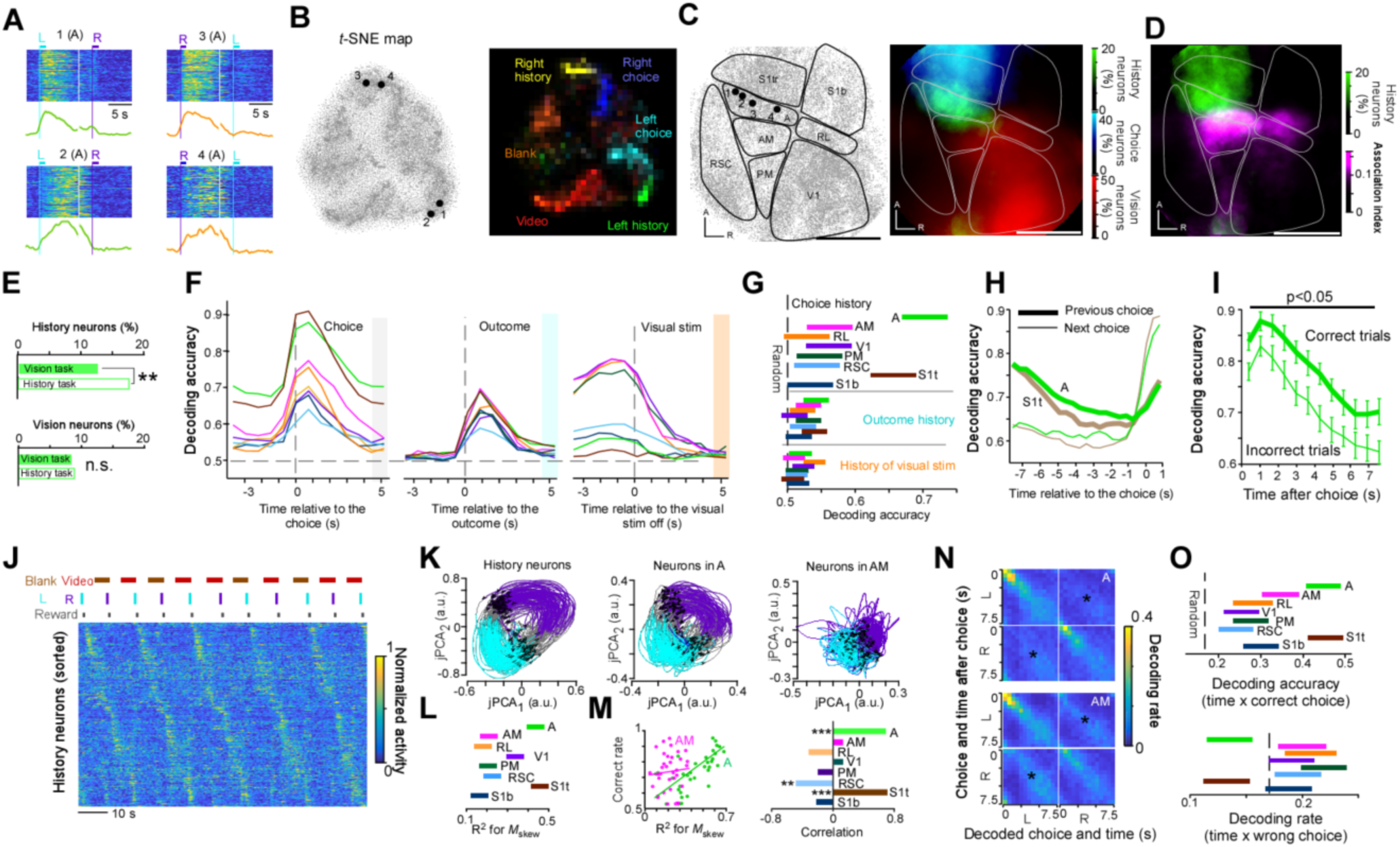
Robust representation of choice history in A. **A.** The representative history neurons. Numbers correspond to that of panel **B** and **C.** Light blue lines indicate rewards delivered from the left lick port, and purple lines indicate rewards delivered from the right lick port. Vertical white lines mark the onset of the sampling period. **B.** The *t*-SNE result for the single neuron activity (left). The right panel shows that each cluster of *t*-SNE has the distinct functional types defined by the encoding model. **C.** Position of the single neurons imaged in history task (left) and the percentages of history, choice, and vision neurons. **D.** The percentage of history neurons and the association index (as defined in Fig. 2J) were overlaid for comparison. Note that these two roughly corresponded to A and AM, respectively. **E.** The percentage of history neurons in A was higher in the history task than in the vision task (top; p=4.1 × 10^-4^; chi-squared test). The percentage of vision neurons in A was not different (bottom p=0.61; chi-squared test). **F.** Decoding accuracy of choice, outcome, and visual stimuli by the activity of 20 neurons from each area using only correct trials, before and after the choice onset, reward delivery, and the end of the visual stimuli, respectively. Line colors corresponded to the areas shown in panel **G. G.** Decoding accuracy of the choice (_p=2.2 × 10_^-13^_;_ one-way _ANOVA_), outcome (_p=0.13;_ one-way _ANOVA_), and visual stimuli (_p=0.094;_ one-way _ANOVA_) at the shaded time in panel **F** were compared for each area. **H.** Decoding accuracy of the previous choice (thick lines) and the next choice (thin lines) as a function of time relative to the next choice. Note that both A and S1t had larger information on the previous choice until 1 s before starting the next choice than the next choice. **I.** The decoding accuracy of the next choice from A activity was significantly larger when the next choice was correct than when the next choice was incorrect (Wilcoxon signed-rank test with Bonferroni correction). **J.** The activity of simultaneously imaged history neurons was aligned by their peak time. The continuous sequential activity can be observed in the two consecutive trials. **K.** jPCA plot of the history neurons shown in panel **J** (left), that of A neurons (middle), and that of AM neurons (right). **L.** The R^2^ of M_skew_ indicating how rotational dynamics was dominant was compared (_p=5.1 × 10_^-33^_;_ one-way A_NOVA_). **M.** The R^2^ of M_skew_ was significantly positively correlated in A and S1t but not in AM (*** p<10^-6^; ** p<0.01; *t*-test). **N.** The decoded rate of time after the choice using neurons in A (top) and AM (bottom). Note that cross-choice mistakes (*) were rarer in A than in AM. **O.** Decoding accuracy of time with the correct choice (top; p=5.6 × 10^-14^; one-way ANOVA) and decoding rate of time with an incorrect choice (bottom; p=8.5 × 10^-10^; one-way ANOVA). Error bars, mean ± s.e.m. in **i**, 95% confidence interval in **G. M,** and **O**. Scale bar: 1mm.

PPC neurons are known to store the information of not only the choice history but also the outcome and sensory histories ^11,13^. We compared these candidates by decoding analysis (**Figure 3F,G**). Although the outcome and visual information rapidly decreased in all areas after a trial, the choice information was strongly represented in A and S1t, and stored during the ITI, especially in A. We confirmed that the choice history information was about the past rather than the future in both A and S1t (**Figure 3H**). We also asked whether the history neurons in A contributed to the working memory. When the next trials were correct, the decoding accuracy from history neurons in A during the ITI stayed higher than when the next trials were incorrect (**Figure 3I**). Thus, the choice history information possessed by history neurons led the animal to the correct choice in following trials. These results suggest that the history neurons exhibited properties of working memory.

What kind of dynamics did the history neurons use to store this choice history information? The population activity of the history neurons indicated that each history neuron had a specific time of peak activity, resulting in the sequential activity after the choice (**Figure 3J**). In this task, left and right selections are alternated so the activity of the history neuron is a sequence that repeats in two consecutive trials. We used jPCA^49^ to visualize and quantify this activity pattern (**Figure 3K**). jPCA identifies low-dimensional projections of population activity that maximize rotational dynamics across time. As expected, the population of history neurons showed one-round rotational dynamics in two trials, and the same was true for the A population, but the AM population did not show rotational dynamics. We quantified the extent to which population activity can be approximated by the rotational component (R^2^ of M_skew_; **materials and methods**) and found that the value in A and S1t was not only higher than that in the other areas (**Figure 3L**) but also significantly correlated with the correct rate of the session (**Figure 3M**). Interestingly, the sequential activity was also observed in AM, but it was less specific to the choice (**Figure 3N**), resulting in the decoding accuracy of time after a specific choice being lower than in A (**Figure 3O**). These results indicate that the history-related activities of A and S1t encode past behavior histories by creating long sequences, and that these rotational dynamics lead to behaviors that reflect past histories.

### Representation of posture

Neuronal activity in S1 typically represents simple somatosensory information, and thus the history neurons in S1t may be a consequence of habitual changes of the posture of the animal after making a choice in a trial. In this case, history neurons may not store the choice history internally, but they may simply reflect the posture of the mouse ^14^. To test this possibility, we analyzed the posture of the mice performing the tasks using a deep-learning based tracking method (**Figure 4A**). We found that the angle of the tail was significantly different from the baseline values several seconds after the response period (RP) (**Figure S5A**). Therefore, the position of the tail was useful information for the next choice in the history task. Overall, although the body was tightly restricted in the body-holder, mice still moved the tail and the paws in relation to the choice, the choice history, and the choice timing. Some parameters could potentially explain apparent task-related activities of neurons, which had been largely ignored in previous studies.

**Figure 4.**
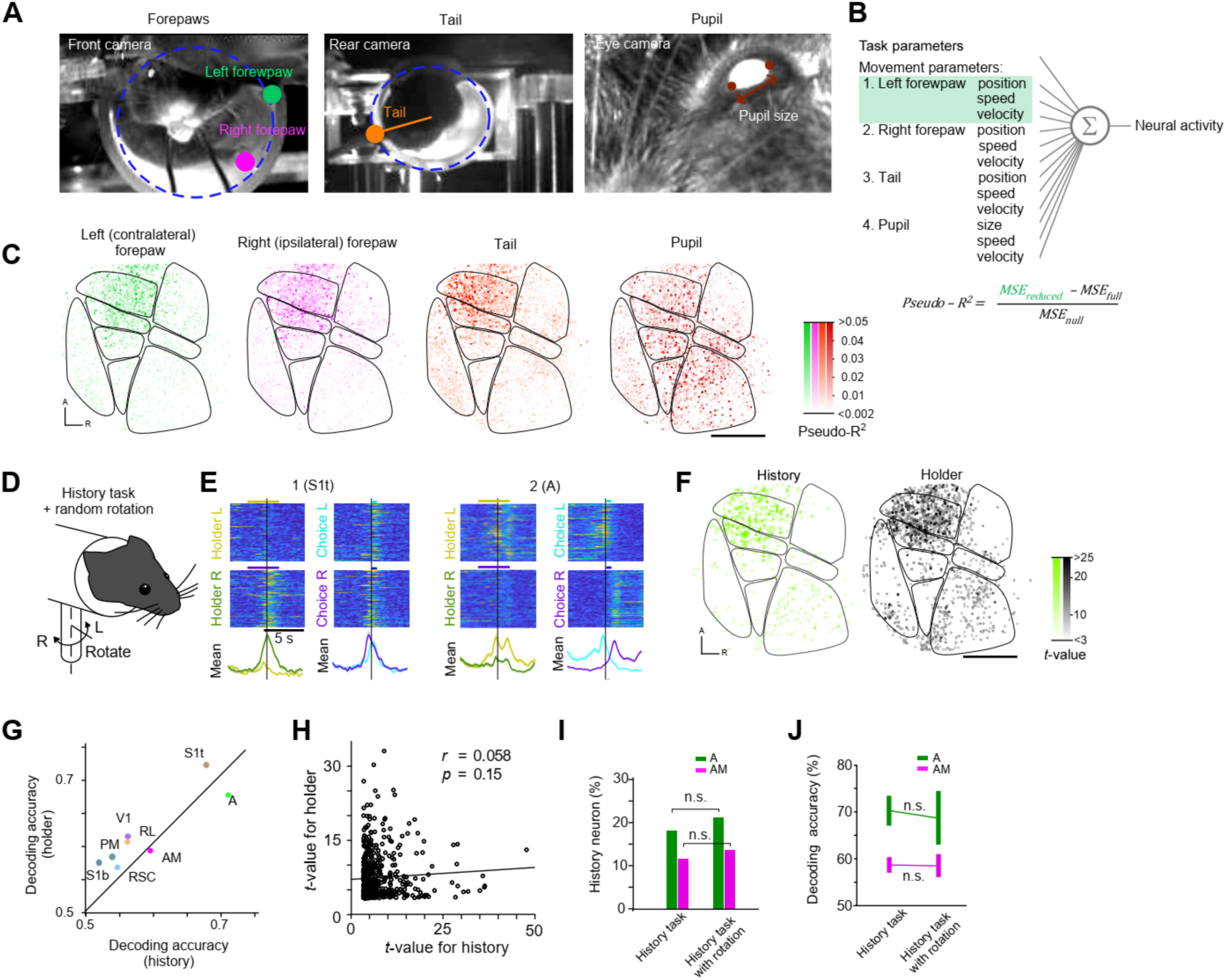
**Posture-related neurons and robust representation of the history neurons. A.** Snapshots from the cameras. The green, magenta, and orange circles indicate that the estimated positions of the left forepaw, the right forepaw, and the tail, respectively, when each part crossed the edge of the body holder. Each line was drawn between each estimated body part and the center of the body holder. Blue circles indicate the edge of the body holder. **B**. Schematic of the regression model for estimation of posture/movement related activity. **C**. Pseudo-R^2^ values were mapped for the left forepaw, right forepaw, the tail and the pupil. **D.** The body holder was randomly rotated during the history task to prevent the mice from keeping choice history as posture. **E**. Representative holder angle-related neurons. Numbers correspond to their location in the left panel. Neuron 1 was holder-related but not to reward direction, whereas neuron 2 was related to both the holder and the reward. **F**. The *t*-values of history neurons and holder-related neurons were mapped. **G**. Decoding accuracy for the holder direction and the choice history in each area was compared. Note that the A and S1t had precise information for both. **H**. The *t*-values for the history-related activity and for the holder angle in each history neuron were plotted. No significant relationship was observed. **I**. The percentage of the history neurons in A (green) and AM (magenta) was not different between the history task and the task with random holder rotation (p=0.15 in A; p=0.38 in AM; chi-squared test). **J**. The decoding accuracy of choice history in A (green) and AM (magenta) was not significantly different regardless of the holder movements (p=0.87 in A; p=0.95 in AM; Wilcoxon’s rank sum test). Scale bar: 1mm.

We searched for relationships between the activity of neurons and these posture-related parameters during the history task. We constructed a linear model based on these parameters to fit the activity of each neuron during the history task in a similar way to the encoding model for defining task-related neurons (**Figure 4B, materials and methods**). We found that the neurons related to the tail and forepaws were similarly distributed around the parietal cortex including S1 and A, while the pupil-size related neurons were mapped around visual areas (**Figure 4C**). Changes in pupil diameter may influence neuronal activity through multiple mechanisms, including behavioral state or noradrenergic level, nonlinear interactions with visual stimulation, and changes in the amount of light reaching the retina. A and S1 had a comparable ratio of neurons that were related to the tail and the paws. However, it remained unclear whether the activity of history neurons represented an internally stored choice history variable, or just reflected the sensory information that happened to correlate with the choice on the next trial.

If neurons in A retain history information by receiving posture information, perturbing the posture information should decrease A’s history information. To test this, we prevented the mouse from maintaining its posture during the ITI by randomly rotating the body holder (+24 or −24 degrees), forcibly introducing a physical disturbance making it difficult for the mouse to retain information of the previous choice by posture (**Figure 4D**). Three mice successfully performed the task in this new condition. We conducted a regression analysis and mapped the neurons reflecting the angle of the holder (“**holder neurons**”; **Figure4F**) and found that they had similar distributions to history neurons (n=3 mice). Decoding analysis also showed the close relationship between history and posture information at the area level (**Figure 4G**). By contrast, the representation of the angle and representation of the choice history were not correlated at the single neuron level (**Figure 4H**). This result is evidence that the neurons regarding two different spatial features, the direction of choice and angle of the body, may share some neuronal resources in A and S1t, but these neurons process the information independently. Consistently, the percentage of history neurons and decoding accuracy of choice history were almost the same regardless of the holder movement in both A and AM (**Figure 4I,J**). Thus, history neurons represent an internally stored choice history variable, independent of the posture. Taken together, the history task and posture-perturbation experiments demonstrated that the most specific function of neurons in area A is storing choice history that contributes to the next choice.

### Mesoscale correlation structure

PPC has anatomical connections with a variety of surrounding and remote areas ^22^, but the functional correlation between those areas has not been extensively examined. In particular, based on the critical functional differences between A and AM that we found, A and AM may belong to distinct cortical networks that consist of different sets of densely interacting cortical areas. To address this question, we focused on the correlation in trial-to-trial variability, *r_t_*. In the following, we analyzed *r_t_* on two different levels: pairs of single-neuron activities and pairs of population activities.

First, to obtain *r_t_* between a pair of single neurons, *r_t_single_,* we averaged the activity of individual neurons over the sampling period by trial type, and subtracted this from the trial-wise activity. The trial types consisted of four sets of pairs of stimuli and responses, that is, the video stimulation and left choice, the video stimulation and right choice, the black screen and left choice, and the black screen and right choice. This operation extracted the fluctuation components of single-neuron activity that are independent of the trial types. Correlation of these fluctuations was defined as *r_t_single._* The average value of *r_t_single_* was small (∼0.01) but larger than 0 even when the distance between neurons was more than 2 mm (**Figure 5A**). Notably, a small number of pairs had a large positive correlation irrespective of distance (**Figure 5B**). In fact, kurtosis (0 in the normal distribution) of correlation after Fisher’s z-transformation was a very large value of 3.77 (p < 10^-6^). Thus, we focused on neuron pairs with *r_t_single_* > 0.3 (shaded areas in **Fig5B**) that were very rare in shuffled pairs in the following analysis to capture the dominant correlation structures.

**Figure 5.**
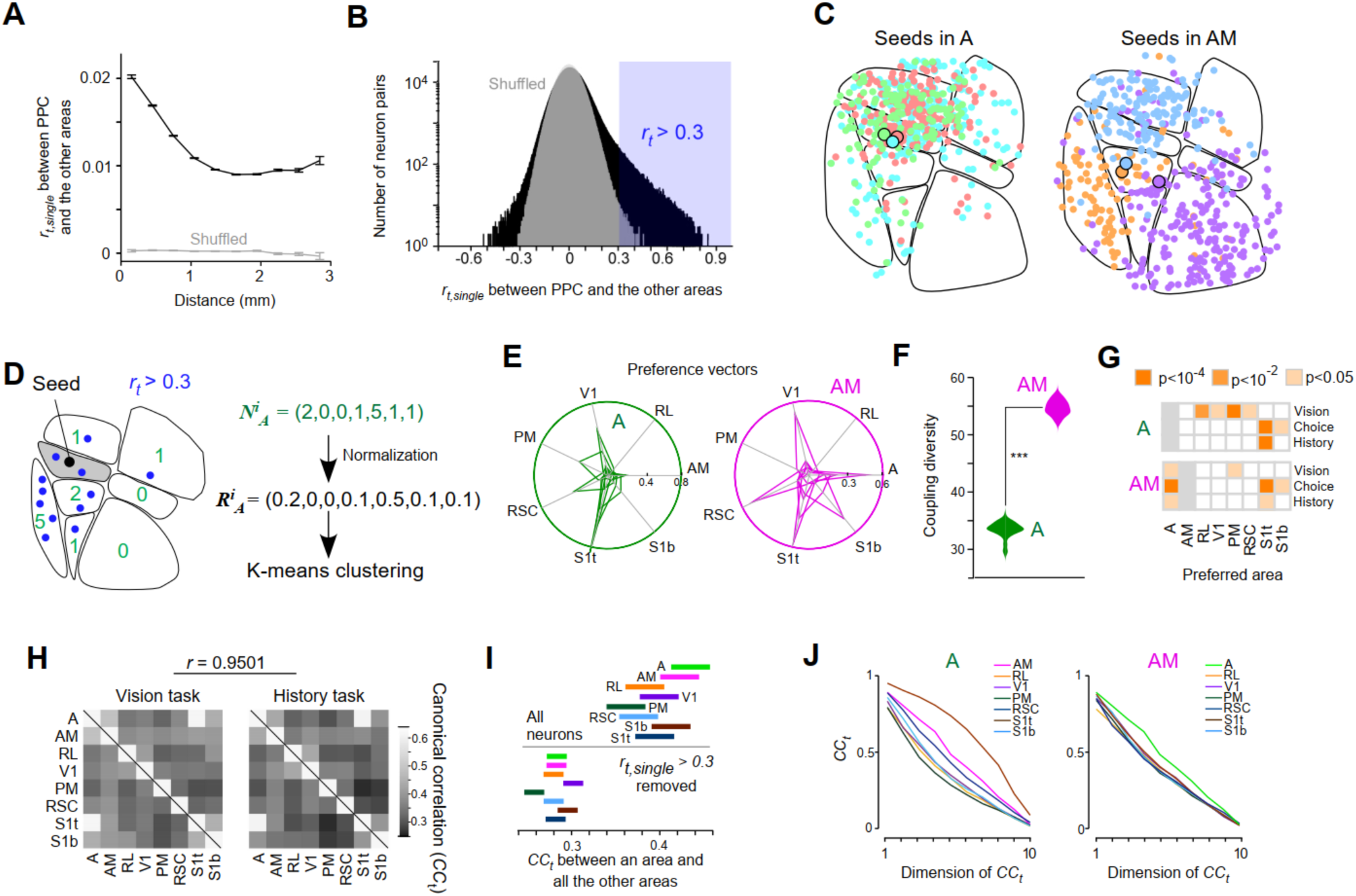
**Mesoscale correlation of PPC and surrounding areas**. **A**. Average of trial-to-trial correlation (*rt*_single) between a PPC neuron and a neuron in the other areas was plotted against the distance between the neuron pair. The correlation was small but significantly larger than zero regardless of the distances. **B.** Log-scaled histogram of the *rt*_single. The shaded area indicates a large positive correlation (> 0.3). **C**. The coupling neurons whose *rt*_single with seed neurons (large circle) in A (left) and AM (right) was larger than 0.3 were shown with the same color as the seed neuron. Most neurons coupling to the A neurons were in the S1t, while the neurons coupling with the AM neurons were spreading but the spreading area was specific to each seed neuron. **D.** Schematic of the K-means (k=6) clustering of neurons in A based on the ratio vector, ***R***. This vector, given for each neuron, consists of the ratio of the number of coupling neurons (*rt_single* >0.3) in the seven areas other than A. In this case, for example, *R^i^A*, the vector of the *i*-th neuron in A, had the largest value of 0.5 in the fifth element, which indicates this neuron had half the number of coupling neurons in the RSP. K-means clustering of the ratio vectors can identify the clusters of A neurons sharing the distribution of the highly correlated neurons. The analysis was done for all neurons in AM instead of A as well. **E.** Polar plots of the preference vectors of A (left) and AM (right). The preference vector is mean ratio vector of each cluster. **F.** Angles between any pair of the preference vectors which were obtained 1,000 times with randomly selected neurons. **G.** The relationship between the preferred area of single neurons in A or AM and their task-relatedness was compared. For example, significantly large number of A neurons preferentially coupling with the S1t neurons were choice neurons or history neurons whereas those coupling with the AM neurons were vision neurons. **H.** *CCt* between pairs of eight areas during vision task and history task was highly correlated (r=0.95). **I.** The *CCt* between an area to the other areas were compared. A and AM had the two largest *CCt* (top; p=1.7 ×10^-6^; one-way ANOVA), which was not the case when coupling neurons were eliminated (bottom; p=6.7 × 10^-7^; one-way ANOVA). **J.** *CCt* as a function of its dimension. *CCt* from AM were uniform regardless of the interacting areas whereas A had large *CCt* with S1t. Error bars, 95% confidence interval.

In which regions are neurons tightly functionally coupled to neurons in AM and A? When mapping the ”coupling” neurons in surrounding areas possessing *r_t_single_* values > 0.3 with randomly chosen ”seed” AM or A neurons, consistent spatial patterns emerged (**“coupling neurons”, Figure S6A**). The coupling neurons were often localized in specific cortical regions. The localized region of the coupling neurons was S1t in most cases when the seed neurons were in A, while the region was variable when the seed neurons were in AM (**Figure 5C**). To quantify this property, we used the normalized spatial distribution of coupling neurons for each individual seed neuron of A or AM (**Figure 5D**). Then we clustered these vectors using K-means (k=6) clustering and displayed the six mean vectors (“**preference vectors**”) for each cluster. The result showed that interaction with most clusters from seed area A was largely dominated by S1t, while each cluster in AM preferred a specific area such as V1, RSC, A, and S1t (**Figure 5E**). These relationships were observed in both vision and history tasks. Thus, AM had multiple interaction channels with surrounding areas, while A interacted with S1t, regardless of tasks. We quantified the similarities of preference vectors by averaging the angles between all pairs of the two vectors, which we referred to as “**coupling diversity**”. We confirmed that the difference in the coupling diversity between A and AM was statistically significant by repeating K-means clustering with randomly chosen neurons in A or AM (**Figure 5F**; **materials and methods**).

Is there a relationship between the interareal interaction and functional types in the task? We termed the area where the largest number of coupling neurons are located the “**preferred area**” and examined the relationship between the preferred area and task relevance. We found that both A and AM neurons with S1t in their preferred areas were significantly more likely to be choice and history neurons. Neurons in area A with visual areas (RL, V1, PM, or RSC) in their preferred areas were significantly more likely to be vision neurons (**Figure 5G**). Moreover, AM neurons preferring A had significant task relevance for all vision, choice, and history. Thus, A and AM neurons determined their functional types by interareal coupling at least in part.

Next, to investigate *r_t_* of the population activity (*r_t_population_*), we first reduced the dimension of population activity in each area into 10 by using PCA (principal component analysis) (**Figure S6B,C**). Then, “fluctuation activity” was recalculated for each dimension and trial type, analogous to the single-neuron analysis described above, but here representing noise in population-level activation patterns. We applied CCA (canonical correlation analysis) to each pair of areas and obtained an average of 10 canonical correlations (*CC_t_*) as *r_t_population._* CCA identifies pairs of linear combinations of population activity from two areas that maximize their correlation across trials, thereby capturing shared population-level fluctuations. The *CC_t_* structure between areas was similar across task types (**Figure 5H)** indicating that this structure reflects the underlying functional connectivity independent of the task. The *CC_t_* between A and S1t was the largest among all the pairs (**Figure 5H**), whereas when the *CC_t_* was averaged across all connections for each area, A and AM had the largest and second largest *CC_t_*, respectively (**Figure 5I**). The dominance in *CC_t_* in A and AM disappeared when the neurons with *r_t,single_* >0.3 were removed. Notably, the *CC_t_* of AM and the other areas was uniform regardless of the paired areas across all 10 canonical components (**Figure 5J**). Thus, area AM is an integration hub of interareal communication, whereas A simply coupled with S1t, and such correlation structure at the population level critically depends on this subset of neurons.

### Relationship between mesoscale functional correlation and anatomical connections

Next, we asked whether distinct network level interaction of A and AM can be seen in the neuropil activity level observed in the same two-photon calcium imaging data. We examined the correlation of trial-to-trial “fluctuations” in neuropil activity (*r_t_neuropil_*). The fluctuation of the neuropil activity was often localized in a specific area or a set of areas (**Figure 6A; Figure S7A**). Consistently, K-means clustering revealed that the cluster boundaries of the neuropil activity approximately corresponded to the area boundaries (**Figure 6B**). *r_t_neuropil_* was similar regardless of the task types (**Figure 6C**) so we combined these data for the following analysis. The neuropil activity in AM well correlated with PM, RSC, V1, RL, and A, whereas that of A highly correlated with S1t (**Figure 6D**). The neuropil activity is known to reflect the local average of input in the imaged location ^50^. Indeed, the trial-to-trial variation of a neuropil activity could be approximated by the average of 1,000 −10,000 neurons within several hundred micrometers from the center (**Figure S7B-D**). These data indicate that the correlation structures of average activity of neurons differ between A and AM.

**Figure 6.**
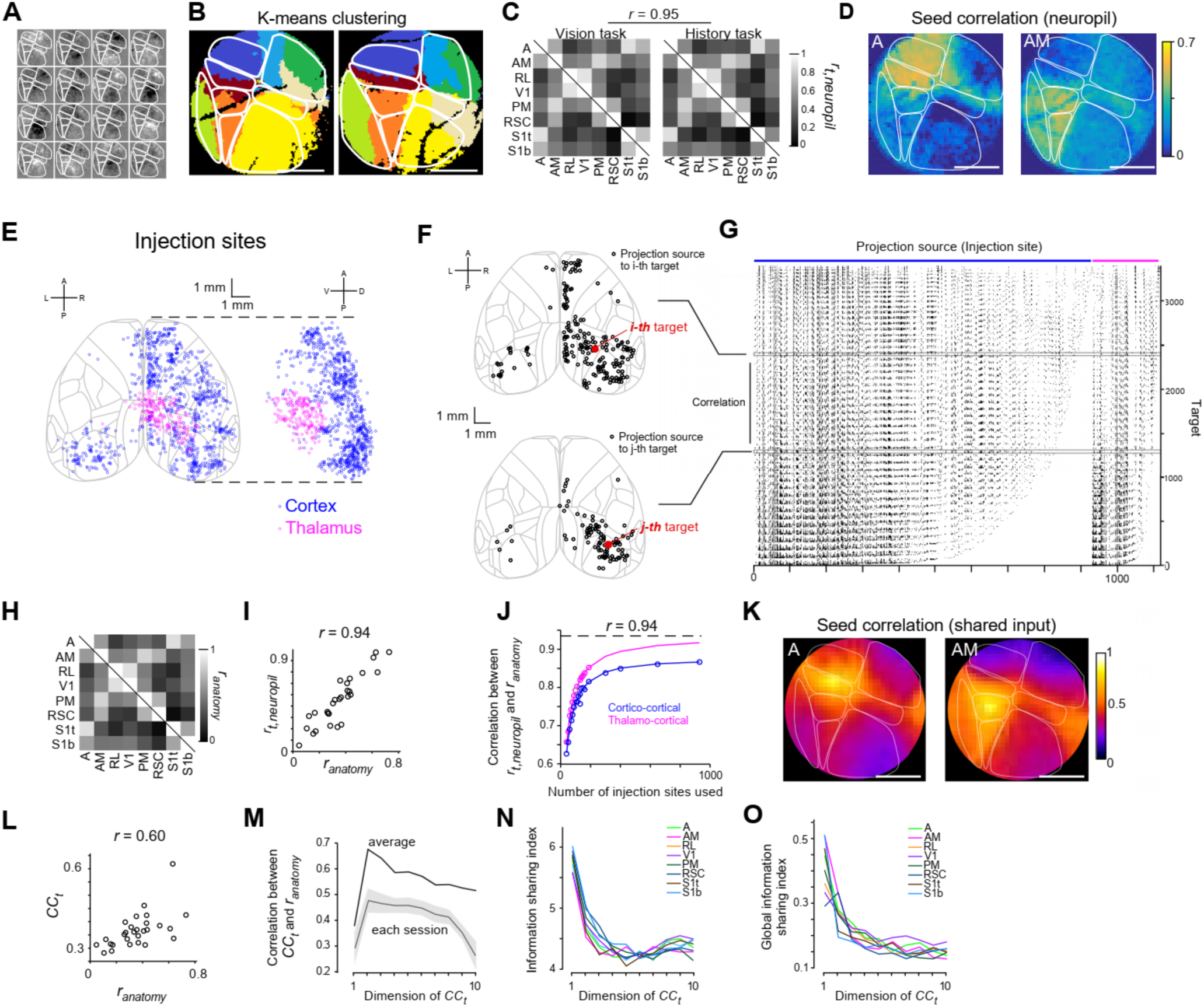
**Comparison of functional and anatomical correlation structures. A.** Trial-to-trial fluctuation of neuropil activity of the mouse 2. Sixteen trials presenting the natural video and when the left choice was made were shown. **B.** K-means clustering of each background pixels (k=8) from two mice resembling the functionally defined areas was shown. **C.** Correlation between background trial-to-trial activity (*rt,neuropil*) in pairs of areas was very similar regardless of the task types (*r*=0.95). **D.** Seed correlation analysis of neuropil activity for A and AM. **E.** Injection sites for cortico-cortical projections (blue) and thalamo-cortical projections (magenta). **F.** Two examples of projection sources (black circles) for a target site (red circle). **G.** A binary matrix Y (*Npix × Nin*j). Each column consists of the projection targets of each injection experiment. Two rows indicated by rectangles correspond to the sets of projection sources for the two target sites shown in panel **F. H.** *ranatomy* between each one of the pairs of 8 areas. **I.** *ranatomy* precisely predict *rt,neuropil*. **J.** Correlation between *rt,neuropil* and *ranatomy* with cortico-cortical or thalamo-cortical projection was plotted against the number of injection sites used for the analysis. **K.** *ranatomy* map when the seed is in A and in AM. Note that these maps resemble panel **D**. **L.** Moderate correlation between *ranatomy* and *CCt* (*r*=0.60). **M.** The first (largest) *CCt* component was the most inconsistent with the anatomical data. **N**. Information sharing index was plotted against the dimension of *CCt*. The first component had the largest information sharing index for all areas. **O.** Global information sharing index was plotted against the dimension of *CCt*. The first component had the largest global information sharing index for all areas. Error bars, mean ± s.e.m.

Next, we utilized Allen connectivity atlas where the mesoscale connectivity between cortical and subcortical areas are comprehensively described. We used results of the anterograde tracer experiments where the adeno associated virus that delivers the fluorescent protein was injected in cortex or thalamus (**Figure 6E**). We found that among cortico-cortical interactions only S1t clearly projected to A more than AM, whereas visual cortices (V1, AL, PM) and frontal cortices (orbitofrontal cortex, dorsal and ventral anterior cingulate cortex) preferentially projected to AM rather than to A (**Figure S8A**). These data are consistent with our results that A represented sensorimotor parameters together with the S1t, and that AM is an interaction hub which was revealed by the mesoscale correlation analysis.

How does information about anatomical connections relate to functional activity correlations at mesoscale? In general, the activity correlation between two neurons is influenced by the common input ^51^. In the case of interareal correlations, we hypothesized that the more common input two areas receive, the higher the activity correlation between the two areas. To address this question, we used the Allen connectivity data to quantify the degree of anatomical input similarity between any two regions as a correlation (*r_anatomy_*, **materials and methods; Figure 6E-G**). Interareal correlation between the anatomical input correlation and the neuropil correlation was surprisingly high (*r* = 0.94, **Figure 6H,I**). This correspondence suggests that the mesoscale interarea correlation is determined by the cortico-cortical and thalamo-cortical common input at mesoscale. The high correlation was more evident when both the cortico-cortical and thalamo-cortical projection data were combined (**Fig 6J**). Thus, *r_t_neuropil_*, which reflects correlation of the average activity of neurons is strictly determined by *r_anatomy_*.

Does this explain the difference in correlation structure between A and AM? The seed analysis revealed that A and AM have distinct patterns of anatomical input (**Figure 6K**) and the patterns highly resembled the neuropil correlations seen earlier (**Figure6D**). This surprising similarity between anatomical and functional seed correlation was consistently observed regardless of the seed areas (**Figure S8B**). Taken together, these data suggest that the distinction of interareal communication in A and AM is led by the cortico-cortical and thalamo-cortical inputs.

### Constraints and limits of anatomical connectivity on neuronal population activity

Although we have so far focused on the differences between A and AM, our data provide broader insights into the relationship between anatomical connectivity and neuronal population activity. First, based on **Figure S7** and the considerations above, anatomical input correlations strongly constrain the correlations between local averages of activity across thousands of neurons. We then asked whether this anatomical constraint extends beyond mean activity, and how anatomical input correlations relate to relationships between neuronal population activities (population vectors).

The correlation between *CC_t_* and *r_anatomy_* was moderate (*r* = 0.60, **Figure 6L**). This moderate correlation did not change when the coupling neurons were eliminated (*r* = 0.61). Interestingly, the largest canonical component was the most unpredictable from the anatomical data (**Figure 6M**). Thus, while interareal correlations based on the mean activity of neuronal populations are largely determined by anatomical input correlations, correlations between population vectors contain additional structure that cannot be captured by anatomical input correlations alone.

One possible source of this additional structure is globally shared activity, which may reflect behavior, brain state, or levels of neuromodulators. To evaluate the contribution of global activity on the canonical correlation between areas, we first compared the canonical coefficient vectors (CCV). We found that the first CCV had a similar orientation, regardless of the paired areas (**Figure6N**). This indicates that the largest components of correlated activity in the CCA analysis are globally shared fluctuations. We also directly evaluated the correlated activity components across all 8 areas with generalized canonical correlation analysis. The first CCV also had a similar orientation to the first generalized canonical coefficient vector (GCCV) (**Figure 6O**). These results indicate that the largest canonical component reflects a global correlation across all cortical areas imaged. Such global correlations may be driven by factors beyond cortico-cortical or thalamo-cortical inputs, such as the animal’s behavioral state as we recently characterized (H. Imamura et al., 2025; F. Imamura et al., 2025). We also confirmed the robustness of these results by repeating analyses using only the 40% highly active neurons after denoising with non-negative deconvolution (36828 out of 91397 neurons; **Figure S9**).

## Discussion

In this study, we functionally characterized neurons in areas A and AM, two components of mouse PPC. Functional imaging during a vision-guided task revealed that neurons in area A showed choice-related activity, while neurons in area AM represented both choice and visual information. Imaging during a history-guided task demonstrated that area A robustly represented choice history with choice-specific activity dynamics. Area A also represented the body posture of the mouse, cooperatively with area S1t. Experiments with perturbation of posture suggested that the choice history-related neurons in A and S1t stored a working memory for the upcoming choice. Analysis of the correlation of trial-to-trial variability indicated that, regardless of the tasks, the interactions of neurons in areas A and S1t were always large and stable, while the interactions of neurons in area AM and the surrounding areas were widespread and shared information specific to each area. These results were consistent with the patterns of corticocortical connections, revealed by both functional and anatomical data. Overall, we determined the functional differences between two PPC regions, areas A and AM, and characterized their distinct mesoscale interactions.

It is generally believed that the parietal association cortex evolved largely after the emergence of primates^52^. It has been unclear whether PPC in mice or rats is useful for understanding parietal association cortex in primates including humans. By contrast, there are commonly observed distinctions within the PPC: Area 5 and 7 in cats ^53^, rostral and caudal PPC of ferrets^54^, galagoes ^52^, squirrel monkeys ^55^, and titi monkeys ^56^ are similar in the sense that visual-to-somatosensory gradient can be observed along the posterior to anterior axis. Macaque PPC also has a similar gradient along rostrocaudal direction ^17–19^. In addition, the somatotopic localization clearly observed in the somatosensory cortex is less clear in area 5 (most anterior part of PPC), where neurons responsive to multiple joints and anterior-posterior limb-tail interactions appear ^44^, similar to responses of area A neurons (**Figure 1G**). Thus, functional differentiation within PPC commonly observed in carnivores, prosimians, New-World monkeys and Old-World monkey was also evident in a rodent. This homology increases the significance of examining the functional differentiation in the mouse PPC in understanding the circuit principle of mammalian parietal association cortex.

The functional differences along the rostrocaudal axis within the medial PPC suggest that various other PPC functions reported previously in mice would be mapped somewhere in this region. Neurons representing decision making ^4-9,29,30^, action history ^11,57^, and the posture ^13,15^ would likely be observed primarily within area A rather than area AM, because area A neurons were found to be activated by multiple somatosensory stimuli, choice, choice history, and body posture. On the other hand, in vision-guided tasks, the neurons in AM would have more fundamental roles in the integration of visual information for coordinating behavior, compared to neurons in A^6,29^. In visual navigation tasks, not only A or AM but also S1t would be highly activated ^7,8,10,12,27,28^. Note that RSC had few neurons with selectivity relevant to the tasks in this study, which suggests that selective activity in RSC may emerge only when mice engage in navigation ^28^ or when the mice need to assess the value of an action ^58^. When vision is critical for choice during navigation and/or determining the value of an action, the neurons in A, AM, and RSC may cooperatively interact ^28^. Fine functional segregation could also depend on task difficulty and learning stage ^8^. However, even when the apparent functional representations are mixed, it is essential to precisely map the PPC subregions to see whether the mixed representation can be decomposed into several groups, each of which is mapped in a discrete subregion.

The differences in forms of interareal interaction among neurons in A and AM imply that AM has a more central role in integrating multimodal information, while A is more specific to controlling and/or monitoring behavior. In fact, area AM interacted with many areas through multiple independent channels, as shown in the correlation analysis, at both the single neuron level and the population level. Neurons in area AM may represent multiple types of information in a category-free manner, and thereby its microcircuit, or downstream circuit, could associate them when needed ^59^. The long-range projections from various cortical areas including frontal cortex, such as the orbitofrontal cortex and anterior cingulate cortex, converge on the AM (**Figure S8A**), supporting the view that AM is a key component in integrative, cognitive roles. Neurons in area A had relatively limited variation in long-range anatomical input, but they had much larger behavior-related activity cooperating with area S1. Thus, our results provide a new framework regarding PPC functions: first, a hub of widespread cortico-cortical interactions (area AM) and, second, a core for behavior monitoring and control (area A) in PPC.

There is another new aspect of PPC function in mice that we present here: the representation of posterior S1 next to the anterior border of PPC. We found that clear somatotopic organization was a critical difference between these two areas. By precisely mapping the borders of PPC and S1, we found that S1t and the anterior part of PPC, area A, possessed almost the same rate of neurons representing choice, choice history and posture, even when posture perturbation was added during the task. Direct functional interaction between S1t and A, and/or top-down input from other areas may support the robust representation of history information in these parietal areas. Area A may send behavior-relevant information to S1 to modulate sensory processing as indicated in the top-down effects from secondary motor and somatosensory areas to S1 ^60,61^. On the other hand, representation of the task-independent posture signals suggests that S1 sends the sensory signals to A. Thus, A and S1 likely send information bidirectionally. These results significantly impact the interpretation of the activity of neurons in the parietal cortex especially during head-fixed navigation tasks ^7-9,27,28^ since the mice always need to coordinate the somatosensory information to maintain body balance during the tasks.

Our data also provided neuropil activity, which may be comparable to the data obtained with widefield, 1-photon calcium imaging that lacks single neuron resolution ^62^. The correlation of trial-to-trial fluctuations in the neuropil activity revealed a clear correspondence to mesoscale anatomical structures. This correspondence was independent of the tasks and was also observed during passive viewing trials (**Figure S8B**). These results imply that shared input from distant areas shapes the spatial structure of cortical activity in general. In other words, the shared input through mesoscale connectivity could predict the brain-wide correlation structures of the functional fluctuations. Since anatomical input correlations can be useful in predicting functional correlations in any cortical area, we developed software that allows for a simple search for seed correlations (**Figure S8C**). We also estimated that neuropil activity approximates the activity of thousands of neurons around the region, suggesting that mesoscale anatomical connectivity constrains the relationship between the activities of two populations of neurons in connected cortical areas. Thus, anatomical structures determine mesoscale functional interactions, and thereby constrain overall activity of local individual neurons. This multiscale interaction would serve as an effective constraint when simulating large-scale neural network models ^63^.

We found that the largest component of canonical correlation reflects globally shared information, which is not well explained by anatomical data. The significant influence of the global component implies that even when understanding the relationship between two areas, it is necessary to consider their relationship to other areas or the entire brain. This global component may involve neuromodulators reflecting states of arousal or motivation ^64^, slow rhythms that propagate throughout the brain ^65^, neuronal activity related to movement ^66^, glial cell activity ^67^, or combinations thereof.

Why are there two different functional modules in the medial PPC, areas AM and A? One possibility is that AM explores potential relationships between various signals from a number of cortical areas and their outcomes. When AM identifies information important to the current situation, neurons in area A store that information for a few seconds and contribute to the action, or behavior. Thus, area AM must constantly compare various pieces of information without bias, whereas area A must focus on specific information. Such a fundamental difference in the required information processing may separate areas AM and A into different functional networks. The coordination of the two would then create a unique and essential role for animals to survive in an ever-changing world.

## Materials and Methods

### Subjects

All procedures conducted at the University of North Carolina at Chapel Hill were reviewed and approved by the Institutional Animal Care and Use Committee (IACUC) of the University of North Carolina at Chapel Hill, which ensures compliance with the standards of the American Association for Laboratory Animal Science (AALAS). All procedures conducted in the University of California, Santa Barbara were carried out in accordance with the guidelines and regulations of the US Department of Health and Human Services and approved by the Institutional Animal Care and Use Committee at the University of California, Santa Barbara. Mice were imaged at the age of 8-40 weeks. Adult C57BL/6J mice (Jackson labs) or GCaMP6s ^47^ expressing transgenic adult mice of both sexes were used in this study. GCaMP6s expression was induced by the triple crossing of TITL-GCaMP6s line (Allen Institute Ai94), Emx1-Cre line (Jackson Labs #005628) and ROSA:LNL:tTA line (Jackson Labs #011008) ^68^. Intrinsic signal of equal numbers of males and females C57BL/6J mice was imaged for the P60 and P100 groups (6 mice per time point, 3 of each sex for each time point) when the variability of the visual areas was examined.

### Surgical procedure and experimental steps for intrinsic imaging

On the day of imaging, anesthesia was induced with 5% isoflurane. After initial anesthesia induction, 2.5 mg/kg chlorprothixene was administered via intraperitoneal injection. Isoflurane was reduced and maintained between 1-2.5% during surgery. Ophthalmic ointment (Lacri-lube, Allergan) was applied to both eyes and a heating-pad was used to maintain body temperature. The scalp was then resected, exposing the skull over visual cortex. To register functional maps of visual cortical areas to stereotaxic landmarks, three reference points were labeled with a marker near primary visual cortex (0.8 mm anterior to lambdoid suture, 2.3 mm lateral to the midline). Using digital calipers, three separate measurements were taken from the center of each dot to the midline (mediolateral coordinates) and from the center of each dot to the lambdoid suture (rostrocaudal coordinates). For each reference point, the three measurements were averaged. We used the lambdoid suture instead of bregma as a landmark since the position of the bregma is less reliable for defining the coordinates of the brain ^69–71^. Although it lacks accuracy, one may be able to use the approximate distance between the lambdoid structure and bregma (∼4.0 mm) to convert the coordinates from lambda-based to bregma-based ones.

A head-plate was affixed to the skull. A 3.5-4 mm craniotomy was performed over the visual areas when needed. The glass window with a 3 mm diameter was put onto the cortex, and a tissue adhesive was used to seal the space between the glass and the skull. Then, the mouse was transferred to the intrinsic signal optical imaging (ISOI) system. The ophthalmic ointment was removed from the contralateral eye prior to imaging by manually blinking the eye gently. Isoflurane was maintained between 1-1.5% during imaging. The camera was focused on the skull to acquire an image of the dots relative to the head-plate and major vessels visible through the skull. The camera was then adjusted for ISOI imaging and an initial retinotopy set was acquired through the skull ^41^. After imaging experiments, the mouse was removed from the ISOI rig and given a second dose of 2.5 mg/kg chlorprothixene. The ophthalmic ointment was reapplied to the contralateral eye.

### Intrinsic imaging and sensory stimuli

The brain was illuminated with 700 nm light and imaged with a tandem lens macroscope focused 600 μm into the brain from the vasculature. Images were acquired with a 12-bit CCD camera (Dalsa 1M30), frame grabber and custom software (David Ferster, with in-house modifications by Jeffrey Stirman & R. H.) at 30 frames per second. Visual stimuli were presented to the left eye relative to the imaged hemisphere using a Dell LCD monitor (Dell U2711b, 2560 × 1440 pixels, 60 Hz) sitting 20 cm from the mouse. The monitor was tilted towards the mouse 17.5° from vertical to cover the visual field (110° by 75°). All stimuli were generated and presented using MATLAB and Psychtoolbox (http://psychtoolbox.org/) ^72,73^. All stimuli were modified to correct for visual distortions caused by the flatness of the monitor. Retinotopy was initially mapped by showing the animal a single, square-wave white bar drifting across a black background to identify right V1 and right HVAs (horizontally for azimuth and vertically for elevation). This stimulus was presented for 30 – 50 eight-second-long cycles.

For somatosensory stimuli, the vibration of the membrane of a standard computer speaker was used. The speaker was controlled with the same LabVIEW software. One side of a plastic pipette was attached to the membrane with tape, and the other side of the pipette was attached to a body part (tail, trunk, neck, left ear, left whisker pad, left forelimb and left hindlimb) gently with tape. As a result, the vibration of the speaker membrane was transferred to the body. A 50 Hz pure tone which gives a 50 Hz sinusoidal movement of the membrane for 0.8 sec vibrated the skin of the targeted body part every 8 seconds. Both cutaneous and proprioceptive responses would be obtained by these stimuli. The speaker sound of 50 Hz tone is far outside the mouse hearing range ^74^, and we confirmed that the stimulation did not evoke activity in the auditory areas with ISOI mapping. The room light was turned off when a pinna or a whisker pad was stimulated to prevent the pipette from being seen since the vibrating pipette was just in front of the left eye. The pipette was outside the visual field from the mouse eyes when the tail, trunk, neck, left forelimb and left hind limb were stimulated.

### Analysis of ISOI data

The images with 12-bit pixel data were binned in software four times temporally and 2 × 2 spatially, resulting in images with 16-bit pixel data. From these binned images, Fourier analysis of each pixel ’ s time course was used to extract the magnitude and phase of signal modulation at the stimulus frequency for mapping of visually evoked areas. The phase of signal modulation was used to generate maps of the phase of the cortical response, mapping retinotopy of the visual cortex. To identify the areas activated by somatosensory stimuli, the mean intensity of each pixel for 2,000 ms after the stimulus onset was subtracted from the mean intensity of the pixel for 1,000 ms before the stimulus onset across all stimulus trials. The negative value of the pixel implies evoked activity in the pixel. The image with evoked activity was filtered with a Gaussian filter with a standard deviation of 10 pixels, and the pixel with the minimal value was identified as a peak location (downward signal is observed in the activated area). The median value of all pixels was set to the baseline. The mean value of the minimal value and baseline was set as the threshold, and the pixels with less than the threshold were regarded as the evoked area.

### Image registration for ISOI map

Retinotopic maps were drawn (ImageJ/Fiji) ^75–77^ based on the first set of imaging data (obtained through the skull) and then modified as needed with the second set (obtained without the skull) to ensure that boundaries between cortical visual areas were accurately identified. These maps were drawn based on color reversals^20^ and other landmarks present in the retinotopy data. For example, when identifying AL, we look for three stereotyped color reversals to identify the border between AL and LM/LI, AL and V1, and AL and RL. We also used the magnitude maps from these stimuli to ensure that the anterior boundary between AL and non-visual areas is accurately restricted to the area that is visually responsive.

The stereotaxic reference coordinates were first plotted, and then registered to the cortical area location by aligning the plotted coordinates with the reference image (image of the stereotaxic reference coordinates marked on the skull). The reference image was then scaled such that 1 mm in the reference image was the equivalent of 1 mm in the coordinate plot. The reference image was then rotated and moved in X-Y (rigid transform) so that the plotted stereotaxic reference coordinates were in agreement with the marks in the reference image. Using the plotted stereotaxic reference markers, the same image transformations were applied to the retinotopically identified ROIs, producing a set of ROIs registered relative to stereotaxic reference coordinates. This was done for each mouse in the data set. The center of mass for each ROI was computed, and this was then converted to millimeters by measuring the number of pixels in 1 mm and dividing the center location by the conversion factor.

### Functional definitions of cortical areas

Previous studies on the demarcation of HVAs based on anatomical tracers have shown that the HVAs comprised AL, RL, A, AM, PM, LM, LI and posterior areas ^20^. With functional mapping with ISOI, we and other laboratories have identified comparable areas with the definition based on anatomical tracing, but the definition of area A and RL has been inconsistent between studies. In this paper, we simply followed our previous studies ^38,39,78^ on the definition of HVAs (AL, RL, AM, PM, LM, and LI) for HVA mapping (**Figure S1**) with ISOI.

In the analysis with two-photon imaging, since we found that area A largely lacked retinotopy, we redefined the areas as follows. By data from two-photon calcium imaging (tactile stimuli and visual stimuli), first, the V1, AM, S1t, and S1barrel were determined by retinotopic and somatotopic organizations. When the cortical neurons activated by visual and tactile stimuli were overlaid, we found an area where the evoked activity was almost absent in the medial space of the FOV. The area was determined as RSC. Then, the triangle behind AM between RSC and V1 was determined as PM. Then, the rectangular space between the S1 barrel and V1, and between S1t and AM was determined as RL and A, respectively. We found a borderline between S1t and S1 barrel, a borderline between V1 and AM, and a borderline between V1 and PM lined up in the straight line. We used the line for the border between RL and A. These definitions are largely consistent with the previous anatomical definitions of the Allen Mouse Brain Common Coordinate Framework (CCFv3) ^79^. We did not discriminate the RSPagl and RSPd in this study since we did not find any functional landmarks. When we analyzed the differences or interactions between areas, we selected the neurons in each area excluding the neurons in a 50 μm-wide transition zone, to avoid contamination.

In addition to our definition, previous studies have also defined posterior parietal cortex (PPC) to include the higher visual areas A, AM, and RL^24^. These areas partially overlap with the parietal association regions defined in the Paxinos atlas, including MPtA, LPtA, PtPD, and PtPR. For a detailed discussion of the correspondence and variability among these regional definitions, see Lyamzin and Benucci (2019) ^24^.

### Surgical procedure for head-fixed behavior

Mice were anesthetized with isoflurane (1.5 – 1.8%) and acepromazine (1.5 – 1.8 mg/kg body weight) when performing a craniotomy over the posterior parietal cortex. Carprofen (5 mg/kg body weight) was administered prior to surgery. Mouse body temperature was maintained using physically activated heat packs during surgery. The mouse’s eyes were kept moist with ointment during surgery. The scalp overlying the dorsal cortex was removed, and a custom head-fixing imaging chamber (^80^; **Figure S3B**) was mounted to the skull with a self-cure dental adhesive system (Super bond, L-Type Radiopaque; Sun Medical) and luting cement (Micron Luting; PD). The surface of the intact skull was subsequently coated with clear acrylic dental resin (Super bond, Polymer clear; Sun Medical) to prevent drying. Mice were then returned to their cages.

After the behavior training and before the two-photon imaging sessions, a second surgery was done with the same anesthetic conditions described above. A 4-mm diameter craniotomy was performed over the posterior parietal cortex (right hemisphere, centered 2.0 mm lateral and 2.50 mm posterior from bregma) and covered with two #1 thickness coverslips with 3.5 mm diameter and one #1 thickness coverslip with 5 mm diameter, glued together with optical adhesive (NOA68, Norland). The edge of the upper coverslip with 5 mm diameter was sealed with luting cement (Micron Luting; PD). Mice were then returned to their cages. The condition of the cortical surface was typically the worst in the first week but recovered after the second week. We did not image the activity of neurons when the cortical surface above the PPC was not clear.

### Head-fixed tasks

In all tasks, mice were kept in a body-holder with the head fixed to the stage (**Figure S3A**). A white square in the left or right position at the display was presented when the mouse licked to the left or right. In the visually guided task (vision task or “V” task, **Figure 2A-C**), mice had to lick left or right based on the visual stimuli presented in the display located at the left visual field. When a natural video taken in the cage (**Figure S3B**) was presented, the mouse had to lick left after a 4-s sampling period to get a drop of water as a reward (1.5 - 2.0 μl) from the left lick port. When a black screen was shown on the display, the mice had to lick right to get the reward from the right lick port. During the sampling period, the lick-ports were withdrawn. Mice learned to not lick during this sampling period (**Figure S3E,F**). The visual stimuli were also presented during the response period and for 2 s after the response when the response was correct. The visual stimuli stopped when the response was incorrect. During the inter-trial-interval (ITI), a gray screen was presented. In the vision task, the probability that the correct direction was left was #R / (#L + #R), where #L and #R are the total number of correct choices in left and right trials before the trial in the session, respectively. When the mice were rewarded three times in the same direction, the correct direction was switched on the next trial. These procedures were implemented to avoid biased licking in the same direction.

We also designed a choice history-dependent task (history task or “H” task) with the same set up as the vision task. In this task, the mice had to choose the other side of the choice from the previous trial (**Figure 3A**). Thus, the information of direction of the previous choice had to be stored for 6-10 s from the previous choice to the next choice including ITI (2 – 6 s) and a sampling period (4 s). Since the visual stimuli were randomly presented during the sampling period, the mice had to ignore the visual stimuli.

In some sessions of the choice history-dependent task, the body-holder was moved randomly to prevent the mice from storing information about the previous trials based on the body-angle. In every trial, the body-holder was reset to the center position when the ITI started and then moved to either a left or right angle (± 24 degrees).

For training sessions before the vision task and history task, we trained the mice with the “V+H” task, where the mice could follow either the visual stimuli and history of the choice. In this task, the correct chosen direction was the opposite of the previous choice as in the history task, and the visual stimuli were consistent with the correct direction. In a few other sessions of history tasks, the display was switched off or the body-holder was randomly moved in the dark environment. We conducted but did not analyze the data of two-photon imaging during the V+H task, the history task in the dark, or the history task in the dark with body-holder movements in this study.

### Training of the tasks

We trained the mice with a serial schedule in custom-made training boxes or under the microscope (**Figure S3C**). After recovery from the surgery of a headplate attachment to the skull, we started water restriction (1 ml / day) until the weight was 85 – 90% of the original weight. Then, we started to train the mice with “pre-training”. On the first day of pre-training, the mice freely got a drop of water from both left and right licking ports with the head-fixed condition for 30 min. During the pre-training after the second session, the temporal pattern of the task (i.e. SP (sampling period), RP (response period) and ITI (inter-trial-interval)) was introduced as the vision task described above except that the RP was unlimited. In other words, the incorrect licking did not result in the termination of RP so that the mice could try to lick until they licked the correct direction and get a drop of water (1.5 - 2.0 μl). The visual stimuli were switched at every trial between a natural video and a black screen, and the correct licking directions were left and right, respectively as in the V+H task. The distance between the two licking ports was set relatively small (∼2 – 4 mm) to allow the mice to find both spouts easily and was gradually increased up to ∼5.5 mm (distance depended on the mice). The experimenter assisted the mice to find the licking port when the mice missed it for several minutes by repositioning it and manual delivery of water. Overall, the pre-training was used to make the mice learn to lick both sides of the licking ports, which took 2 – 5 sessions. When the number of rewards the mice received exceeded 150 per session (< 2 hours), the task was switched to the V+H task. In this task, the temporal pattern of the task was the same as the vision task, but the correct direction of licking was set to the opposite side of the previous direction. Therefore, the mice could either follow visual stimulus or the choice history to get the reward. The experimenter assisted the mice to find the licking port by repositioning the spouts and manual delivery of water when the mice lost motivation, especially in several initial sessions. The V+H task was trained to give the baseline condition where we can retrain the V task and H task from the task. After the acquisition of the V+H task, we switched the task to the vision task. The correct rate immediately dropped after switching from V+H task to V task but recovered after that. The transient decrease in the correct rate indicates that the mice used history information during the V+H task. After reaching the criteria, the mice were trained to perform the task under the microscope. After the V task, we retrained mice with V+H task to recover the baseline condition. After that, we switched the task to the H task. We found that the correct rate dropped immediately after switching the task from V+H task to H task and recovered again. This transient drop demonstrates that the mice also used vision information in the V+H task. In the H task, all the mice reached the criteria under the microscope during two-photon imaging (blue dots in **Figure S3D**). With two mice, we trained the H task with randomly moving body-holder, and both mice reached the criteria during the two-photon imaging (orange dots in **Figure S3D**). Throughout the training, mice kept their weight more than 85 % of the original weight and, and their health conditions were monitored daily for approximately four months.

### Two-photon imaging

Two-photon imaging was carried out using a custom Diesel2p microscope controlled by custom LabVIEW software (National instruments). The two-photon excitation laser from an 80-MHz Newport Spectra-Physics Mai Tai DeepSee was scanned by a resonant scanner (CRS 8 KHz, Cambridge Technologies) and two Galvo scanners (62010H, Cambridge Technologies). The photon signal was collected by a PMT (H7422P-40, Hamamatsu) and amplified with a variable high speed current amplifier (#59-179, Edmund Optics). Two-photon imaging of 3 × 3 mm^2^ was collected at 5.6 Hz for imaging. Since the maximum scan angle of the resonant scanner is limited, we made two rectangles with 1.5 × 3 mm^2^ to cover 3 × 3 mm^2^ FOV (**Figure S2A,B**). With these parameters, a neuron with a 10 μm diameter circle was supposed to have 29 pixels inside the ROI (region of interest), including more than 100 laser pulses on average, which was sufficient to find the single neuron activities **(Figure S2C,D)**. Imaging was performed with <120 mW excitation 910 nm laser out of the front of the custom-made objective lens (0.55 NA). Mice were head-fixed at about 15 cm from a flat monitor, with their left eye facing the display during imaging. An 80° × 45° region of the left visual field was covered with the display. During imaging, we monitored a diameter and position of the left pupil, movements of whiskers, both forelimbs, a mouth and a tail with four custom-controlled CMOS cameras at 30 fps. Licking to the left or right licking port was also monitored with an electrical circuit ^81^. The timing signals for the resonant scanner were used for synchronization of all the other devices and the data. The imaging depth below the dura was between 80 μm and 180 μm (layer 2/3). To ensure the scanning planes were parallel between different sessions, we tilted the stage of the body holder manually before the imaging sessions to make the imaging plane horizontal to the surface of the brain.

Visual stimulation for retinotopic mapping was displayed on a 60 Hz 7” LCD monitor (9.5 x 15 cm^2^). A white vertical or horizontal bar was drifted from temporal to nasal or top to bottom at a cycle of 8 s. The tactile stimulation for mapping of the somatotopic organization was made with a gentle and brief air-puff (20 – 50 ms) to a specified part of the body at a cycle of 4 or 8 s. Both visual and tactile stimuli were repeated at least 20 cycles. When the mouse started to actively move during the stimuli, the imaging was stopped and the data were discarded.

### Image processing

Image analysis was performed using ImageJ (National Institutes of Health) and MATLAB (MathWorks, Natick, MA, USA) software. Image sequences were corrected for focal plane displacement, and region of interest (ROIs) within somata were automatically determined using suite2p **(Figure S2E)** ^82^. The fluorescence traces and shape of ROIs were inspected visually. Elongated ROIs, which were more likely to be dendrites or axons, were excluded if the ratio of the length of the major axis to the length of the minor axis was greater than **3**. To eliminate the effect of background activity, we made a square-shaped mask whose pixels were within 50 μm from the edge of the ROI, excluding pixels less than 5 pixels from the edge of the ROI and pixels within the other ROIs. Then, we subtracted the mean value of this area from the mean value of the ROI. The obtained value was referred to as “activity” of the neuron. Those neurons showing activity with skewness of > 0.3 ^80^ were regarded as **“active neurons”** and used for analysis. We also obtained active neurons with kurtosis > 5 and defined them as **“highly active neurons”**. Non-negative deconvolution was conducted with foopsi ^83^. The values after non-negative deconvolution were thresholded with median + 1.5 × 1.4826 × mad and was shown as the inferred spike.

To analyze neuropil activity, the pixels within each ROI identified with suite2p were filled with each background activity described above. After that, the size of the image was reduced to 120 × 120 pixel^2^, where the size of a single pixel is 25 × 25 μm^2^.

### Processing of video data using DeepLabCut

We recorded video data (160 x 120 pixel^2^, 30 fps, ∼ 1 hour / session) from three IR cameras (Basler SCA640-70GM, GIGE, Lens: Basler C125-1218-5M, 12 mm) placed in front of the mouse, at the rear of the mouse, and in front of the left eye during task performance. We analyzed these data with DeepLabCut ^84^ (https://github.com/AlexEMG/DeepLabCut) for tracking the left and right forepaws, the tail and the pupil. We labeled each body part in randomly extracted 100 frames from each video and trained a model for each video data for each session using the 100 labeled data. About 20,000 iterations were performed in training the model. Google Colaboratory was utilized for the training of the model using GPU. We further analyzed the positions of the body parts in frames where the estimated relative likelihood was greater than 0.9. The horizontal and vertical position was directly extracted from the result of the tracking, and then we obtained the direction (angle) of the left and right forepaws and the tail from the body center (**Figure 4A**). The pupil size was calculated by identifying two edges of the left pupil (**Figure 4A**). We visually inspected the quality of the results and confirmed that all the tracking results were as precise as human vision.

### Encoding models

To map the neurons encoding visual, choice and history-related information, we used a simple linear encoding model (**Figure 2G**). We confirmed that the range of maximum VIF (variance inflation factor) for parameters for visual, choice, and history was 1.7 - 3.5 (mean: 2.5) in the vision task, 1.5 - 3.1 (mean: 2.0) in the history task, and 2.1 - 4.1 (mean: 2.9) in the history task with holder rotation. In total, the range was 1.5 - 4.1 (mean: 2.4), which indicates that simple linear regression is sufficient to estimate the correlation between these parameters and single neuron activity without a serious problem of collinearity. The activity of each neuron was fitted by a set of predictor variables for each session. The 29 - 32 predictors consisted of the timing of the visual stimuli (556 ms each, 11 bins, video (+1) or dark (−1)), chosen directions (left (+1) or right (−1)), choice histories (3.7 - 7.4 s after left (+1) or right (−1) choice), reward directions (2 s after rewarded, left (+1) or right (−1)), rotation of the body holder (left (+1) or right (−1)), and an inter-trial interval. Non-selective parameters were also added for each parameter, to avoid identifying neurons activated by both types of stimuli, choice, history, reward or holder rotation as task-related neurons. For example, the non-selective parameter was +1 on the time of choice regardless of the choice directions. The Bonferroni-corrected *p*-values for the visual stimuli, choice, choice history, and holder rotation parameters in the encoding model were used to identify the vision, choice, choice history, and holder-related neurons, respectively. P-values (p<0.001) for each parameter of the regression model were used to determine the significance. The *t*-values in the regression model were used to map the strength and position of each task-related neuron. To map the ratio of task-related neurons, we counted the number of active neurons and the number of task-related neurons within 150 μm of each pixel and divided the latter by the former to calculate the ratio. The association index was determined by the harmonic mean of the rates of vision neurons and choice neurons. The harmonic mean approaches the arithmetic mean when the two values are similar, but becomes closer to the smaller value when the two values differ substantially. Therefore, the association index takes a large value when both vision neurons and choice neurons are abundant.

We also mapped the posture/movement parameters by estimating the posture/movement-related neurons with a regression analysis. As some of the VIFs for the predictors including task parameters and posture/movement parameters were very high (typically 300 or more), we used a linear model with L2 parameter penalization (a ridge regression). The predictors consisted of position, speed, and velocity of the left forepaw, the right forepaw, and the tail along the x-y plane and the angle **(Figure S5)**, pupil size, speed and velocity of pupil size change, and task parameters described above (total 59 parameters). The neuronal activity and predictors were z-scored. We determined the lambda value for each neuron (range: 10^-4^ ‒ 10^4^) and avoided overfitting with five-fold cross-validation. To estimate the contribution of parameters for the left forelimb, the right forelimb, the tail, and the pupil, we repeated the same analysis with a reduced model where each set of predictors was eliminated from the full model (**Figure 4B**). Then, the pseudo-R^2^ was obtained for each set of predictors by (*MSE_reduced_* ‒ *MSE_full_*) */ MSE_null_*, where MSE is the mean squared error, *MSE_reduced_* is MSE for the reduced model, *MSE_full_* is the MSE of the full model, and *MSE_null_* is the null model. The null model predicts a fixed value that is independent of the explanatory variables; specifically, it simply outputs the mean of the training data. For example, we constructed a regression model without the parameters regarding the left forelimb (green shade of Figure 4B), obtained *MSE_reduced_* for the left forelimb, and the pseudo-R^2^ was calculated as above by comparing the MSE of the full model and the null model. This value reflects the extent to which the position of the left forelimb contributes to the prediction of neuronal activity.

### Decoding analysis

The direction of the choice was decoded by the population activity of neurons with a logistic regression model (Matlab built-in function, “glmfit” with binomial distribution and logit link function). For each time bin, the model was trained to predict the chosen direction using the activity of neurons in the time bin except for the trial. Then, the choice in the trial was predicted by the activity of neurons in the trial (leave-one-out). The outcome (reward or punishment) and visual stimulation (video or blank) were similarly decoded by the activity of neurons. The number of trials was matched to the smaller number of the right and left selections in order to align the number of selections between the left and right selections. For example, if the left selection had 150 trials and the right selection had 200 trials, all trials from the left selection were used and 150 trials from the right selection were randomly selected. To avoid bias due to the selection of trials, we repeated the random selection 20 times and averaged the 20 decoding results to determine the accuracy of the decoding. By this manipulation, decoding accuracy is 50% when the neuron has no information at all.

The time after the initiation of the videos (**Figure 2L**), or the time after the left or right choice (**Figure 3F,N**) was independently decoded by using a support vector machine (Matlab built-in function, fitcsvm) and the activity of neurons. For decoding the time after the video starts, each imaging frame (185 ms) during the sampling period was used as a label for classification of the time. For decoding the time after the choice, 9.5 seconds after the choice were divided into 17 parts (each part being 555 ms) and each part was used as a label for the classification. We trained multiple support vector machines, each of which classifies whether a given set of neural activities belongs to the *i*th interval or not (*i* = 1, 2, …, the total number of parts). A ten-fold cross-validation procedure was used to avoid over-fitting.

### Dimensional reduction

To confirm whether the encoding model explains the diversity of single neuron activity, we visualized each set of the task-related neurons with *t*-SNE. We averaged single neuron activity over trials for each of four trial types: video + left choice, video + right choice, blank + left choice, and blank + right choice. Then, we defined a vector for each neuron by combining the four averaged activities. The Matlab function ”tsne” was applied to this set of vectors.

jPCA is a dimensional reduction method for visualizing neural dynamics developed by Mark Churchland (https://churchland.zuckermaninstitute.columbia.edu/content/code) ^49^. It is useful for visualizing not only rotational dynamics but also sequential dynamics because it approximates the direction of change in the population vector over time that is orthogonal to the current activity (*M_skew_*). To apply jPCA analysis, the activity of neurons in the area of interest was smoothed with a Gaussian filter with σ=0.9 s and z-scored. After reducing the dimensionality to 6 using standard PCA, the activity from 1 s before to 5 s after the choice was concatenated on all trials. jPCA was applied and a linear transformation of the jPCA in the first plane was used to show the neural trajectories. To compare the neural dynamics with behavioral performance, we applied jPCA to 10-minute segments of population activity and compared R^2^ for *M_skew_* with the correct rate in those 10 minutes. R^2^ for *M_skew_* is the coefficient of determination of the regression model for approximating the temporal change of population vector by the current population vector ^49^.

### Correlation analysis at single neuron and population level

Correlation of trial-to-trial variance of activity between a pair of single neurons was defined as *r_t_single_.* To calculate *r_t_single_*, we averaged the activity of individual neurons over the sampling period, and the average across each trial type was subtracted from this value. The trial types consisted of four sets of pairs of stimuli and responses, that is, the video stimulation and left choice, the video stimulation and right choice, the black screen and left choice, and the black screen and right choice. By this operation, we extracted the fluctuating components of single-neuron activity that are independent of the trial types. Although, the finding that neurons with high *r_t_single_* tend to share the functional properties we propose is not a trivial consequence of the analysis. At the same time, it remains possible that high *r_t_single_* reflects the degree to which neurons share unobserved features, and that such features are correlated with our functional classification. Thus, while this analysis suggests that correlated fluctuations across cortical areas may contribute to the determination of functional types, establishing an exclusive conclusion will require more fine-grained behavioral measurements, tighter control of internal states, and causal identification through targeted interventions.

We defined a neuron with 0.3 or more *r_t_single_* with a neuron in AM or A as a coupling neuron if the neuron was not in the same area. We counted the number of coupling neurons in each area and considered the number as a vector. We normalized this vector so that the sum was 1 (**ratio vector**) only when the total number was 5 or more. Then we clustered the ratio vectors using K-means (k=6) clustering and defined the mean vector of each cluster as a “**preference vector”**. We confirmed that repeated K-means clustering with different initial values resulted in that more than 80% of the neuron pairs being classified into the same cluster. To quantify the diversity of coupling patterns across clusters, we computed the angle between every pair of preference vectors. We then averaged these pairwise angles and defined this quantity as the “**coupling diversity**”. For comparison of coupling diversity of AM and A, we randomly extracted half of the ratio vectors from AM and A, conducted K-means clustering, and obtained the coupling diversity 1,000 times to know the distribution of the coupling diversities. We defined area X as the **“preferred area”** of a neuron in A and AM if the neuron had one or more coupling neurons in the area X. We then compared the proportion of task-related neurons (i.e., vision, choice or history neurons) between the neurons whose preferred area includes area X and the other neurons by a chi-square test.

To calculate the trial-to-trial fluctuations in neuropil activity (*r_t_neuropil_*), we averaged the neuropil images over sampling period, and the average image across each trial type was subtracted from this image. The trial types consisted of four sets of pairs of stimuli and responses, that is, the video stimulation and left choice, the video stimulation and right choice, the black screen and left choice, and the black screen and right choice. By this operation, we extracted the fluctuating components of neuropil activity that are independent of the trial types.

### CCA (canonical correlation analysis)

CCA was used to compare neural population activities in two areas. As a preprocessing step, the data from an area was reduced to *N_PCA_* (=10) dimensions using PCA (principal component analysis). This ensured that CCA did not find dimensions of high correlation but low data variance ^85^. To estimate the area-to-area correlation in the task-independent variables, the CCA was calculated in task-independent spaces, which is practically the same as noise correlation, as follows (**Figure S6B,C**).

The value of each principal component was averaged over a 4-s sampling period in each trial. Then, we obtained the fluctuation component of the value by subtracting the mean of each four trial-types. Here four trial-types were defined by a combination of visual stimuli and choices: natural video and left choice, natural video, and right choice, black screen and left choice, and black screen and right choice. We call these fluctuating dimensions task-independent space. The fluctuation was concatenated into one column vector per each principal component (*N_PCA_* by *N_trial_* matrix, *N_trial_* is a total number of four trial types). CCA between any pair of the cortical areas was obtained in these task-independent spaces. The *N_CCA_* (=10) CCA values were averaged and the value was referred to as *CC_t,X,Y_*, where X and Y are a cortical areas.

We also used CCA to estimate the amount of shared information between an area and the other areas (**Figure 6L,M; Figure S6D,E**). Suppose we have time series of the *N_PCA_* variables (principal components) in the task-independent spaces, **X** and **Y_a_**, from an area X and another area Y_a_, respectively,

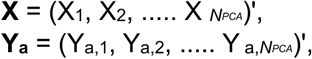

where X*_i_* and Y_a_,*_i_* (*i*=1,2,3,… *N_PCA_*) are column vectors, whose length is equal to the number of trials (*N_trial_*). To calculate *N_CCA_* canonical correlation values, linear transformation of **X** and **Y_a_** with weight matrices (canonical coefficient vectors, CCVs) **A_a_** and **B_a_** give canonical scores, **S_a_** and **T_a_**, respectively as follows:

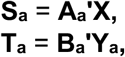

where both **A_a_** and **B_a_** are *N_PCA_* by *N_CCA_* matrices with *N_CCA_* CCVs (column vectors) of length N_PCA_. The *n*-th CCV transforms the time series of principal components, **X** and **Y_a_,** to *n*-th canonical score. As a result, **S_a_** and **T_a_** are the matrices with *N_CCA_* by *N_tria_*_l_. This transformation was done so that the correlations between the canonical scores are maximized (with the built-in function “*canoncorr*” in Matlab).

We have seven time series of the N_PCA_ variables (principal components) in the task-independent spaces, **Y_1_, Y_3_, …, Y_7_**, from areas Y_1_, Y_3_, …, Y_7_. Then, we can get sets of CCVs, **A_2_, A_3_, …, A_7_** and **B_1_, B_3_, …, B_7_** and sets of the canonical scores, **S_1_, S_3_, …, S_7_** and **T_1_, T_3_, …, T_7_.** Here, the CCVs of **A_1_, A_2_, …, A_7_** defined in the 10-dimensional data space of area **X** correspond to the linear projections of activity of neurons in the area X that best explain the activities of neurons in each of the other areas (7 areas in total). If area X shares similar information with another seven areas, CCVs, **A_1_, A_2_, …, A_7_** will be oriented in a similar direction and therefore the length of the sum of the CCVs will be larger. Accordingly, we defined the **“information sharing index”** (**Figure S6D**) of area X as the length of the sum of CCVs as max 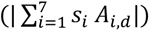, where 𝑠_!_ can be 1 or −1, 𝐴*_i,d_* is a *d*-th column vector (CCV) of *i*-th area, and || denotes the norm.

We also used generalized CCA (GCCA)^86^, which extends the notion of CCA to >2 data sets. We estimated similarity between the first generalized CCV and CCVs for each area by inner product of these vectors. We defined the average of these inner products as **“global information sharing index”** (**Figure S6E**).

### Analysis of Allen connectivity data

We used the Allen Mouse Brain Connectivity Atlas (http://connectivity.brain-map.org ^33^) to investigate the pattern of the corticocortical and thalamocortical projections around the mouse PPC. In this database, connectivity data are available for mice from injection areas (AAV1-EGFP as an anterograde tracer) to any brain region. For estimating the relative projection density between A and AM, the corticocortical projection density was obtained in each layer at 0, 100, 200, 300, 400, 500, 600, 700 μm from pia, averaged and normalized with the injection volume, then averaged across wild-type mice. This was repeated with the major source areas to the A and AM (V1, PM, AL, vRSC, S1t, OFC, dACC, and vACC; **Figure S8A**). For seed correlation analysis, we downloaded the data whose injection sites were either isocortex (929 experiments) or thalamus (189 experiments) at 100 μm resolution (**Figure 6E)**. We set a 100 μm-space grid that covered entire dorsal cortical areas including the PPC. Then, a set of injection sites that projected to each grid at the 300 μm from the surface of the brain were identified for each grid (**Figure 6F**). Thus, each grid has a set of the injection sites where the grid receives its projection from either isocortex or thalamus. Using these binary connectivity matrices, we calculated the correlation coefficient between two sets of injection sites of the two grids. Based on this correlation matrices, we got a map of the correlation coefficient for each grid (seed) (**Figure S8B**).

Let a binary data matrix **P** (*N_pix_* × *N_inj_*) be a set of results of the tracer experiments, where *N_pix_* and *N_inj_* is the number of pixel (cortical surface) and the number of injection experiments, respectively (**Figure 6G**). Let **Q** (*N_pix_* × *N_inj_*) be a set of the injected volume (area), ad let **W** (*N_pix_* × *N_pix_*) be a pixel-to-pixel connection matrix. By definition,

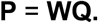

Here, we need a correlation matrix, **WW^T^**. However, there are no direct data for **W** or **Q**. Therefore, we estimated it by **P** as follows

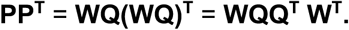

If the injection volume is small and random enough, **QQ^T^** → **Id** as the number of experiments gets large. If so, we can approximate **WW^T^** by **PP^T^**. In reality, **QQ^T^** should work as a spatial filter of **WW^T^**. As the injection volume would be <500 μm on average, the seed correlation map should capture the structure with a resolution of several hundred micrometers, which would be sufficient for our analysis. This resolution could be improved with decreasing injection volume, or kernel regression ^87^.

To effectively visualize these maps, we developed a software that users can explore cortical surface point (seeds) to see the cortical areas that shared either the corticocortical input, thalamocortical input or both types of input (**Figure S8C**, https://web.ece.ucsb.edu/~riichirohira/TopBM-2.05/index.html).

### Statistics

Wilcoxon’s rank-sum test and chi-squared test were used for pairwise comparison. For multiple comparison, we used one-way ANOVA and showed confidence intervals (p=0.05) in the figures. In **Figure 2K** and **N**, and **Figure 3G**, **L, M**, and **O,** the bars indicate the 95% confidence intervals. All other bars denote s.e.m., unless otherwise noted. No statistical tests were performed to predetermine sample size. No blinding or randomization was performed. For the decoding analyses, the number of animals was treated as the independent variable, whereas for the encoding model analyses, the number of neurons was treated as the independent variable. To ensure that the results were not driven by a single animal, we repeated the statistical tests while systematically excluding data from one animal at a time and confirmed that statistical significance was preserved in all cases. Furthermore, qualitative interpretations were made only when differences in effect size were clearly observed.

## Data availability

Data is available upon request.

## Code availability

A web-based interactive software including its source code is available online (https://web.ece.ucsb.edu/~riichirohira/TopBM-2.05/index.html). The script for the data analysis is available upon request.

## Acknowledgements

This work was supported by National Science Foundation (grant nos. NeuroNex 1934288 and 1707287 to S.L.S.; BRAIN EAGER 1450824 to S.L.S.) and the NIH (grant nos. NINDS R01NS091335 and NEI R01EY024294 to S.L.S.; R01NS128079 to I.T.S.; JP22wm0525007 from AMED, to R.H. and JP22H02731 (RH), JP21B304 (RH), JP21H05134 (RH), JP21H05135 (RH) from MEXT/JSPS to R.H.

## Supplementary figures

**Figure S1.**
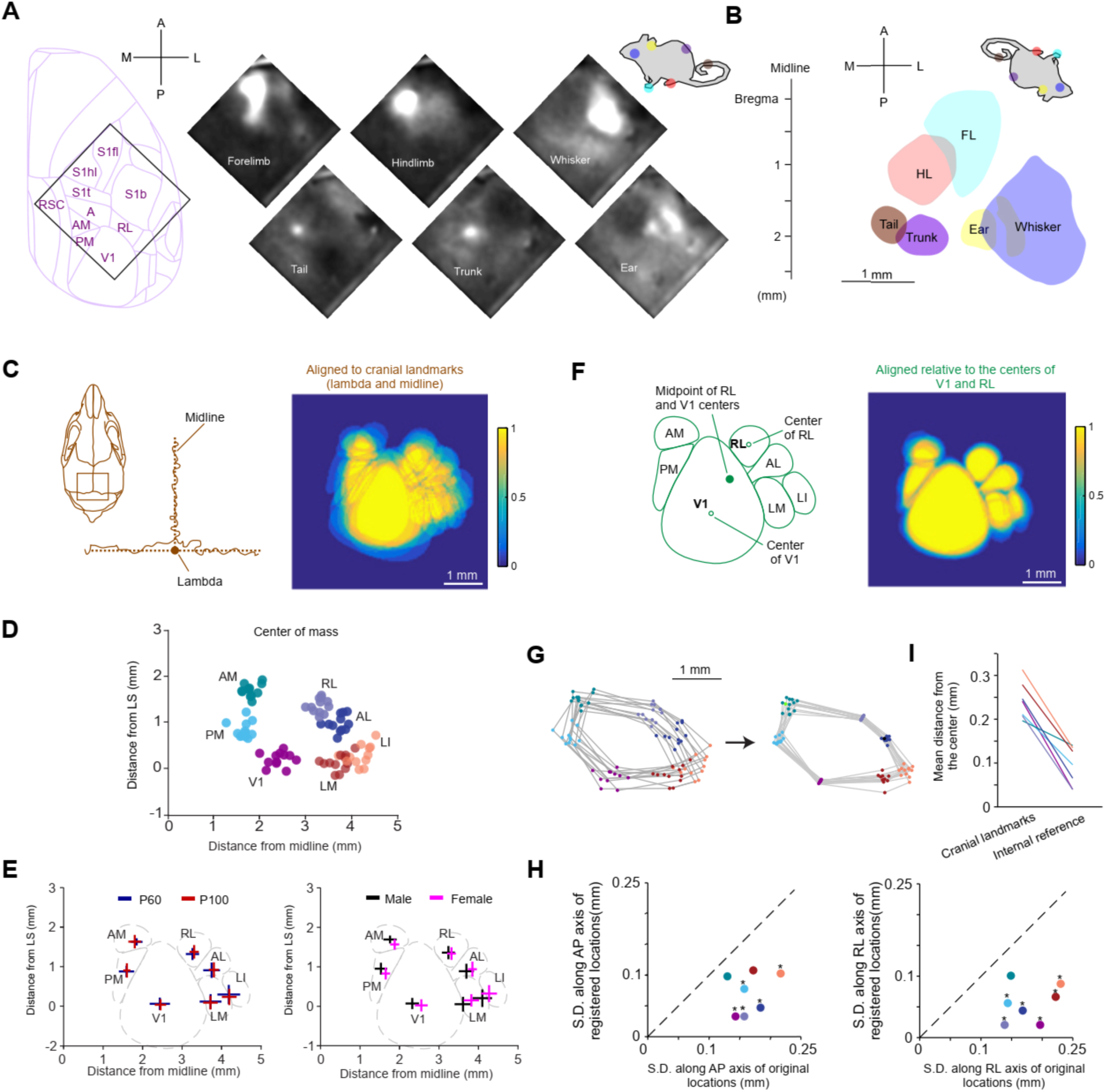
Intrinsic signal imaging and the individual variability of the position of the HVAs. **A.** Activity maps of S1 for forelimb, hindlimb, whiskers, tail, trunk, and ear were shown. **B.** S1 regions obtained in panel **A.** A “homunculus” was shown for reference. **C.** Left, mouse skull and the bony landmarks for reference. We used the midline and a lambda on the skull to register the V1 and HVAs across different mice. Right, overlapped maps of the V1 and HVAs for 12 mice. Note that there is large variability of HVA locations across mice. **D.** Distribution of the center of mass of V1 and HVAs relative to the midline and lambda. **E.** Comparison of the distribution of the center of mass of V1 and HVAs between different ages (left) and between different sexes (right). No significant difference was found (p>0.05, Wilcoxon rank-sum test). The bar indicates ± standard deviation. **F.** Left, we used the midpoint of the center of V1 and the center of RL, and the angle between the midline and the straight line connecting the center of V1 and the center of RL to register the V1 and HVAs across different mice (internal registration). Right, the overlapped maps of the V1 and HVAs for 12 mice. Note that all the HVAs are better registered than the panel **C**, right. **G.** Location of V1 and HVAs before (left) and after (right) the internal registration. The gray lines connect the points from the same mice. The dot colors are the same as panel **D**. **H**. The standard deviation of the location of the center of mass along the anteroposterior and mediolateral axis before and after the internal registration. * p<0.05. ** p<0.01 (Two-sample *F*-test for equal variance). The dot colors are the same as panel **D**. **I**. The mean distance from the center position of each HVA in each mouse to the average center position across mice was compared before and after the registration.

**Figure S2.**
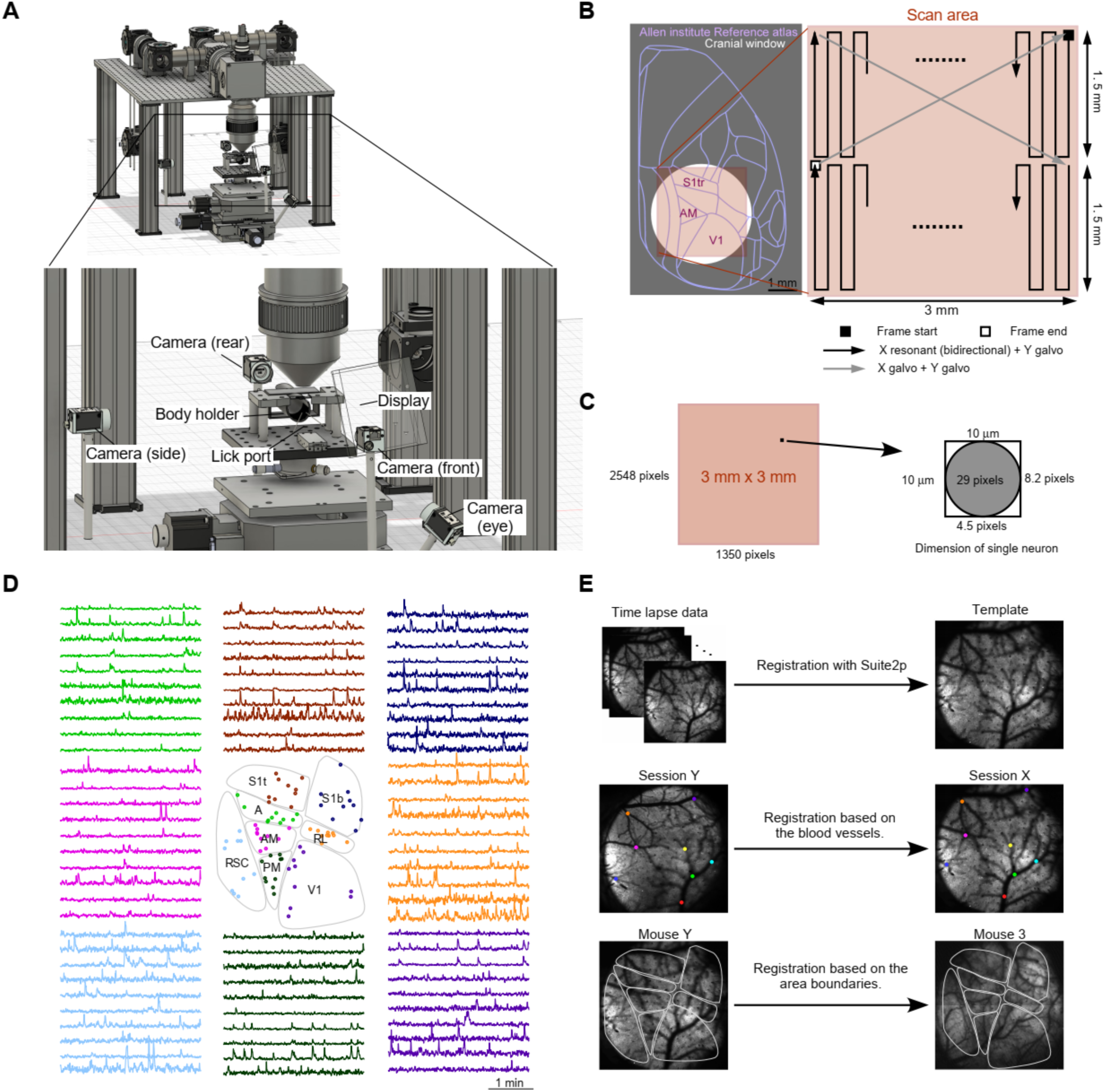
A large field-of-view two-photon microscope and its scanning strategy. **A.** A large field-of-view (FOV) two-photon microscope, Diesel2p ^16^. The body holder, head-plate holder, display and four cameras are also shown. **B.** Left, the brain areas of the right cerebral cortex of the mouse from the Allen atlas (version 3) ^33^. Right, the scanning strategy to cover the 3 mm by 3 mm square area around the mouse PPC. Two 1.5 mm by 3 mm rectangle areas were sequentially scanned using two Galvo scanners and one resonant scanner. **C.** The scanning strategy in panel **B** resulted in 2459 x 1350 pixel^2^ for each square frame, which yields 29 pixels (>100 laser pulses) for a circle with a 10 μm diameter (the size of a single neuron). **D.** Representative activity of simultaneously detected neurons in the same FOV. Ten neurons in 8 areas are shown. Each color of trace corresponds to the area. **E**. Top. The image sequence was registered to the template within a session. Middle. The day-by-day distortion was corrected based on the blood vessels. Bottom. When overlapping the neurons across different mice, we registered images based on the boundaries of areas.

**Figure S3.**
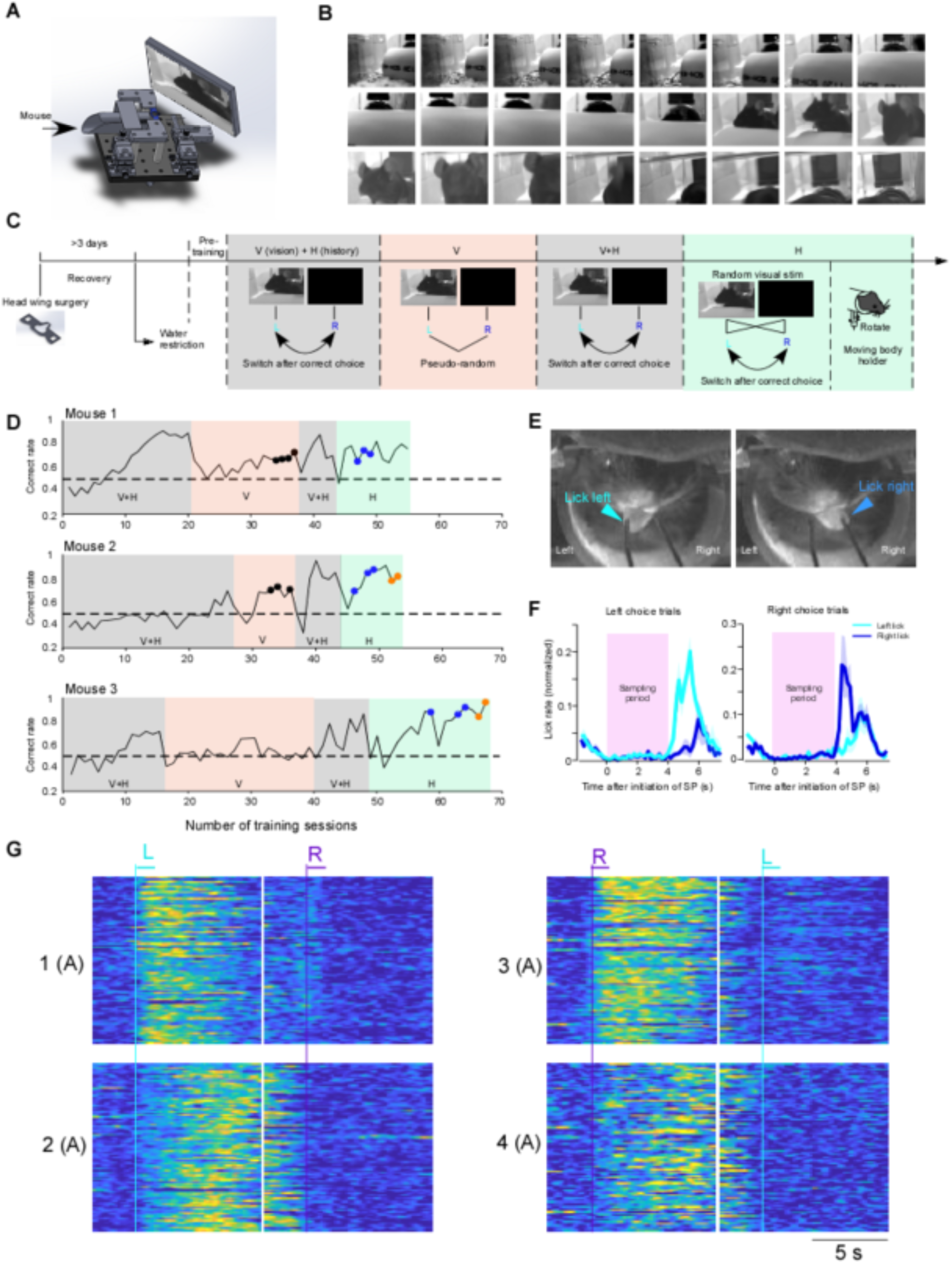
**Training schedule and learning curves of the tasks for head-fixed mice**. **A.** A mouse was head-fixed and held inside a plastic body-holder. The mouse can enter the body-holder smoothly with its head plate into the aperture of the body holder. The display was located at a location covering the left visual field. **B.** A montage of 4-second natural videos (30 Hz, 24 out of 120 images) was shown. This photo was taken while moving a camera in a cage. **C.** The serial training schedule for multiple tasks. **D.** The correct rates of three example mice were plotted as a function of training sessions. Background color corresponds to the type of the task illustrated in panel **C**. The black, blue and orange dots indicate the imaging experiments of the vision task, history task and history task with a moving body-holder**. E.** The tongue extension movement during the task was monitored by the video put in front of the mouse (30 fps). Note that this does not necessarily indicate contact between the tongue and spout. We electronically measured contacts between the tongue and the spout and used the signal for the choice. **F.** Offline analysis captured the tongue extension to the left (left licking, magenta) and tongue extension to the right (right licking, green) during the response period. In both trials with the left choice (top panels) and the right choice (bottom panels), tongue extension movements during the sampling period were hardly observed. **G.** Enlarged view of Figure 3A. All four neurons exhibit multiple calcium transients during the delay period.

**Figure S4.**
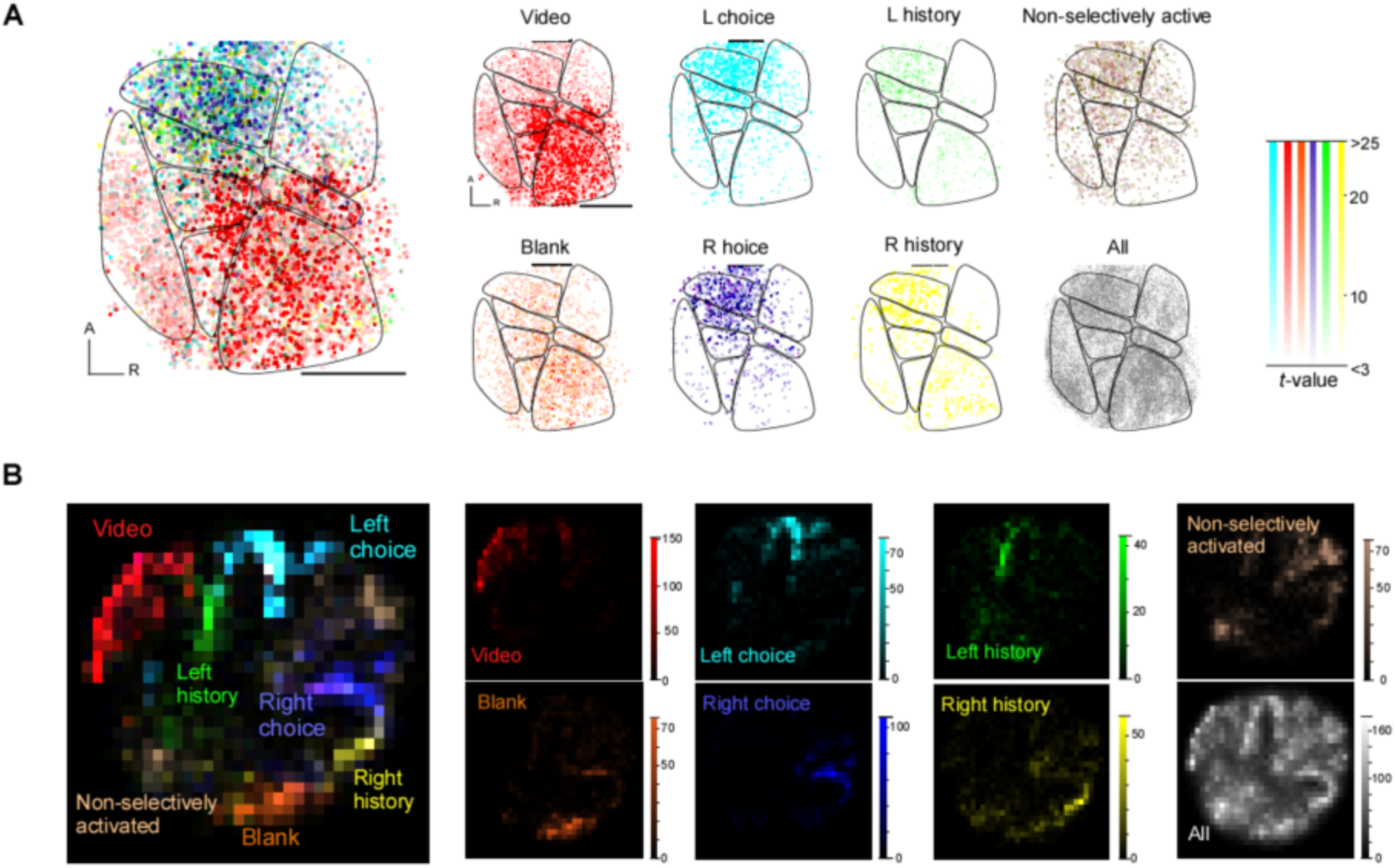
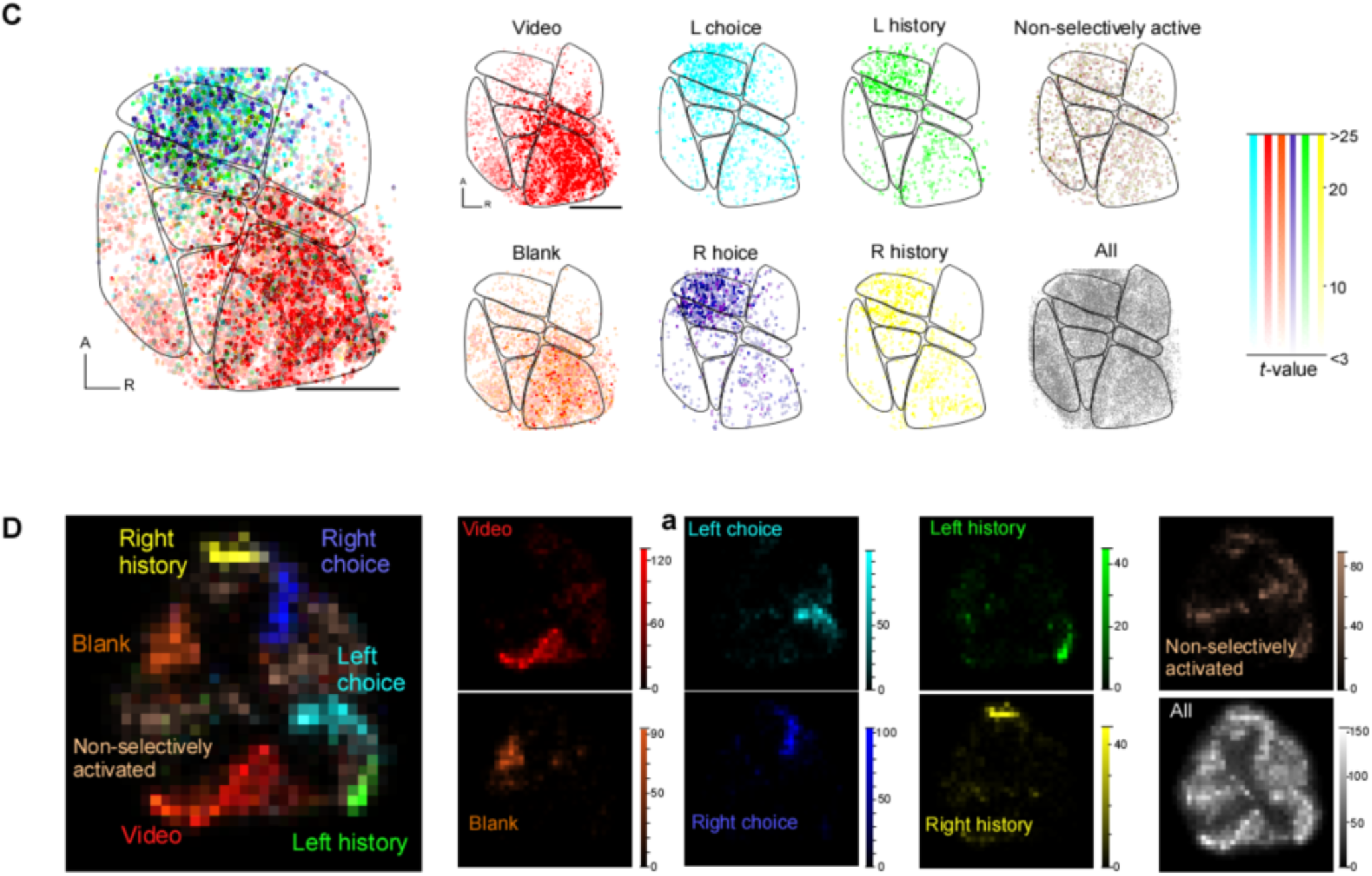
Distribution of the task-related neurons in vision and history task. **A**. The *t*-values for each type of task-related neurons were mapped in the vision task. **B.** The result of *t*-SNE analysis in the vision task. **C** and **D**. The same as **A** and **B** but in the history task, respectively.

**Figure S5.**
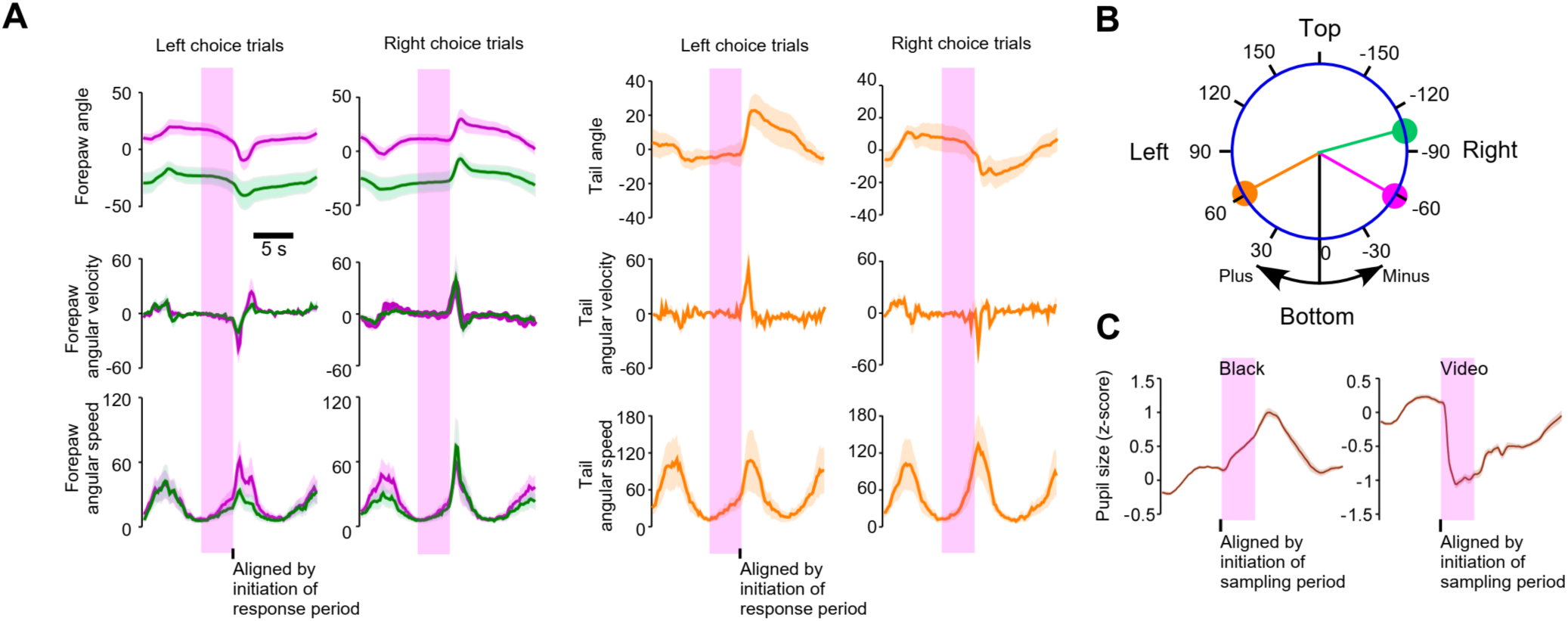
Analysis of postures and movements during the task. **A**. The position, the velocity and the speed of the three body parts were aligned by the initiation of the response period (RP) for the trials with left and right choice during the mice performing the history task (n=3 mice, total 6 sessions). Shaded areas indicate the sampling period (SP). **B.** Coordinates of the angle of the body parts. **C**. The pupil size increased and decreased after the black screen and the natural video started, respectively.

**Figure S6.**
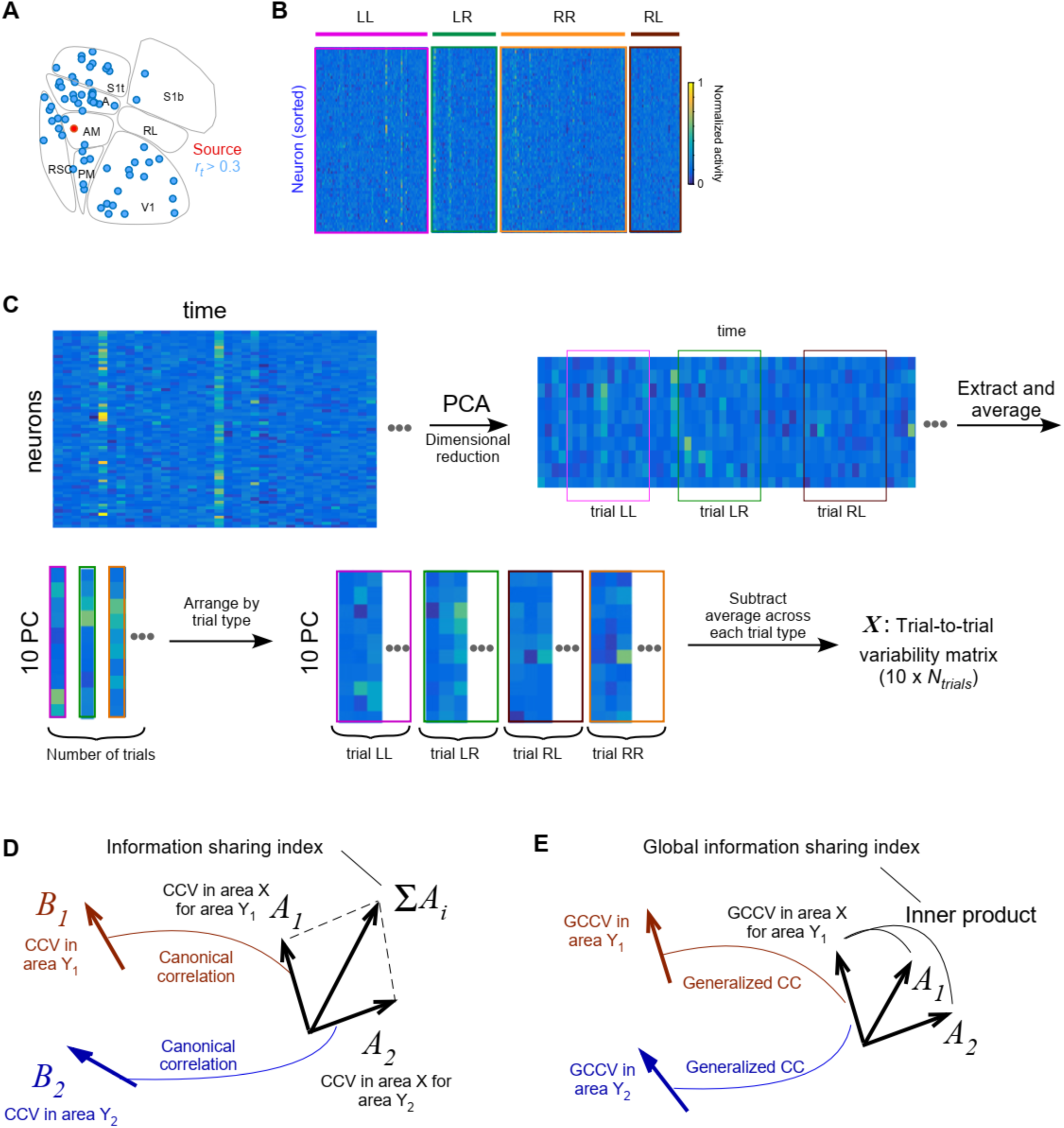
**Canonical correlation analysis (CCA)**. **A**. An example source neuron in AM (red) and neurons with *r_t_* ≥ 0.3 (blue) were shown. **B.** Trial-to-trial activity fluctuation of the highly correlated neurons (*r_t_* ≥ 0.3, panel **B**) was color coded in each trial block. **C**. Illustration of CCA pre-processing. Activity of the population of neurons was extracted in each area, and the PCA (principal component analysis) reduced the data dimensions to 10. Then, “fluctuation” activity was calculated similar to the single neuron case. The data during the sampling period of each trial block (**LL**:video/left choice. **LR**:video/right choice. **RR**: black/right choice. **RL**: black/left choice) was extracted and averaged. Then, the residual of the data after subtracting the trial-mean in each trial type was concatenated as a new matrix, ***M_t_*** (one column vector per PC dimension, that is, 10 × *N_trial_* matrix). **D.** Illustration of the canonical correlation analysis and calculation of information sharing index. See methods for the precise definition. **E.** Illustration of the generalized canonical correlation analysis and calculation of global information sharing index. See methods for the precise definition.

**Figure S7.**
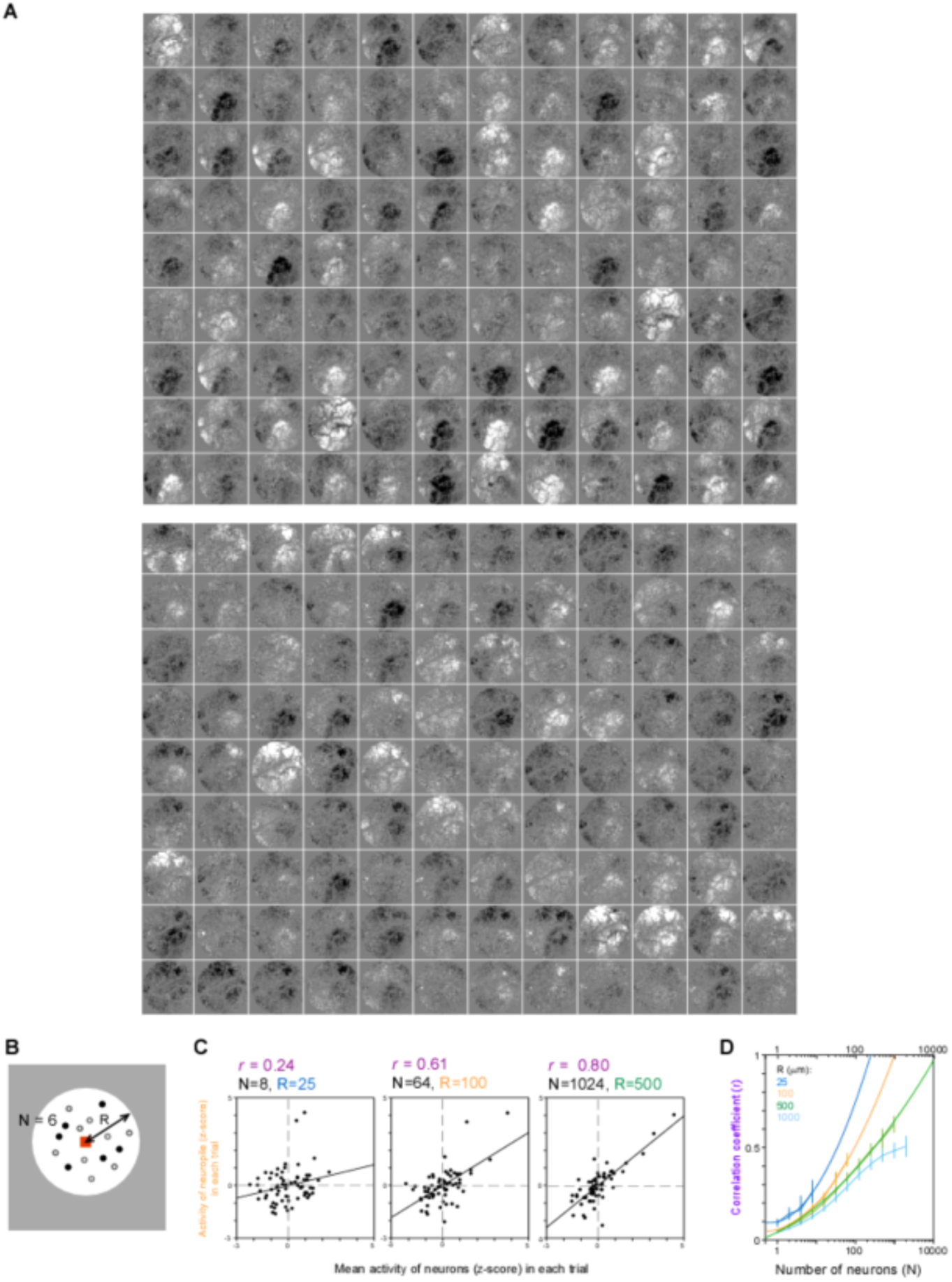
The trial-to-trial fluctuation of the neuropil activity and its relation to the single neuron activity. **A.** The average neuropil activity during the sampling period was shown for each of the 108 trials. The top panel shows data collected during the Vision task, and the bottom panel shows data collected during passive visual stimulation. Boundaries between cortical areas were clearly visible. **B.** To compare the trial-to-trial fluctuation of the neuropil activity and single neuron activity, the single neurons were randomly selected (filled circle) within the distance, R, from the center (red square). Then, the trial-to-trial fluctuation of the neuropil activity at the center and the trial-to-trial fluctuation of the mean activity of selected neurons was compared. **C.** The z-scored neuropil activity in each trial was compared with the z-scored mean activity of *N* selected neurons in the corresponding trial. **D.** The correlation coefficient obtained in the panel **C** was repeatedly calculated with a range of neurons averaged and the distance. The lines show the polynomial fitting curves.

**Figure S8.**
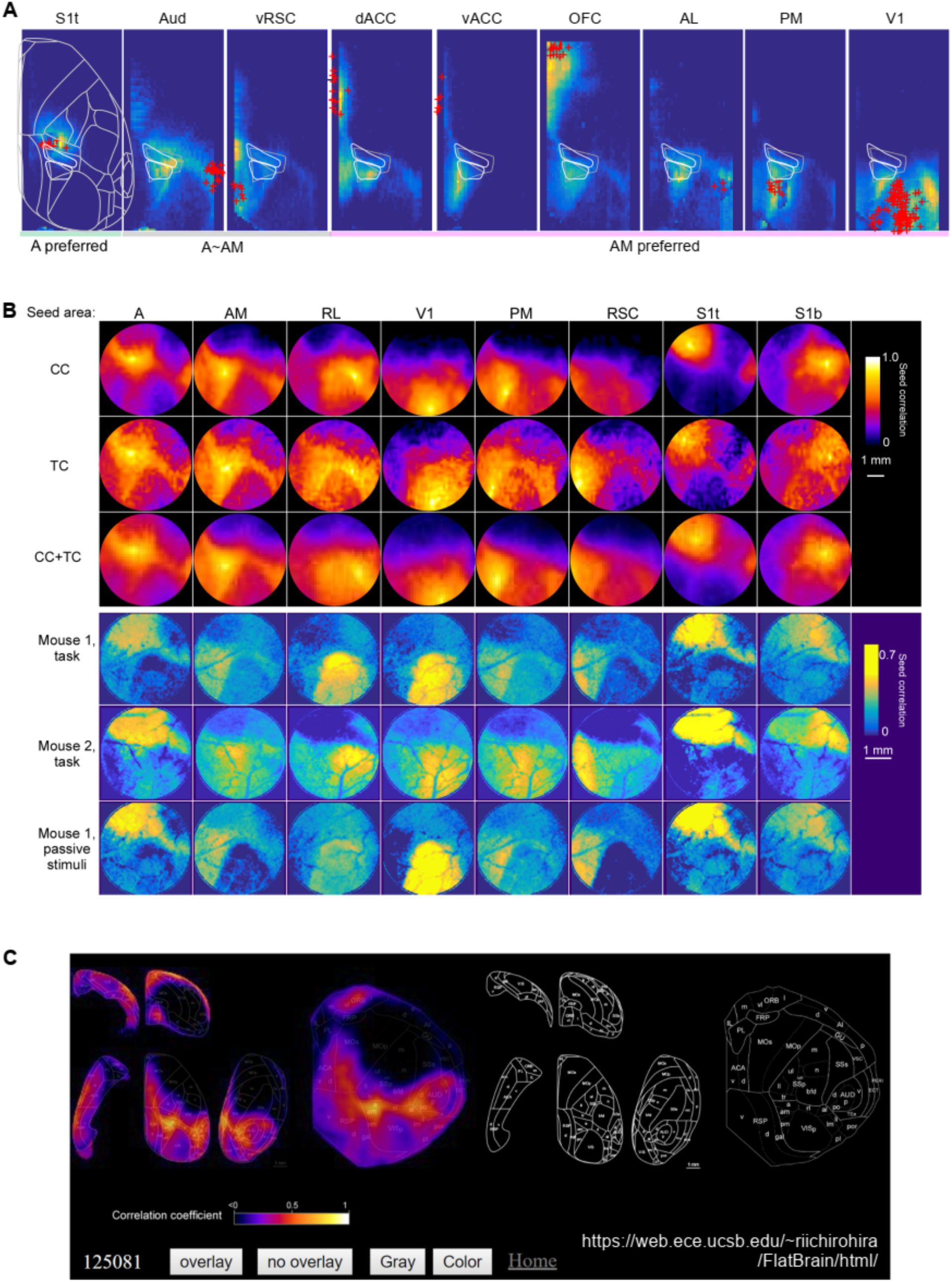
Anatomical structure analyzed with anterograde tracer database and its relation to the correlation structure of the trial-to-trial variability in neuropil activity. **A.** Using Allen connectivity atlas, the axonal density of cortico-cortical and thalamo-cortical projection was analyzed. The nine cortical areas where A or AM receive dense projections are shown. The density was obtained from multiple depths (0, 100, 200, 300, 400, 500, 600, 700 μ m from pia), averaged and normalized with the injection volume, then averaged across the experiments. Only the projection from the S1t was preferentially distributed in A rather than AM. Top-down axons from dACC, vACC, and OFC preferentially projected to AM rather than A, as well as visual cortices (AL, PM, and V1). Gray boundaries show A and AM in Allen brain atlas, and the white boundaries indicate A and AM in our definition. **B.** Top and second rows, the correlation between each cortical region and the seed points in each area (A, AM, RL, V1, PM, RSC, S1t, S1b) was color-mapped. The high correlation indicates that the region has highly shared input from the cortex (“CC”, top row) or the thalamus (“TC”, second row) with the seed region. A dotted circle indicates the imaging area. Second-bottom and bottom, the seed correlation of trial-to-trial activity fluctuation during the vision task (“M2, V”, second-bottom row) and during passively viewing the moving bar (“M2, P”, bottom row) of mouse 2. Note that all the correlation maps in each column are similar. **C.** A snapshot of the online interactive software, “BrainModules”, for exploring the anatomical seed correlation analysis. Link: https://web.ece.ucsb.edu/~riichirohira/TopBM-2.06/index.html

**Figure S9.**
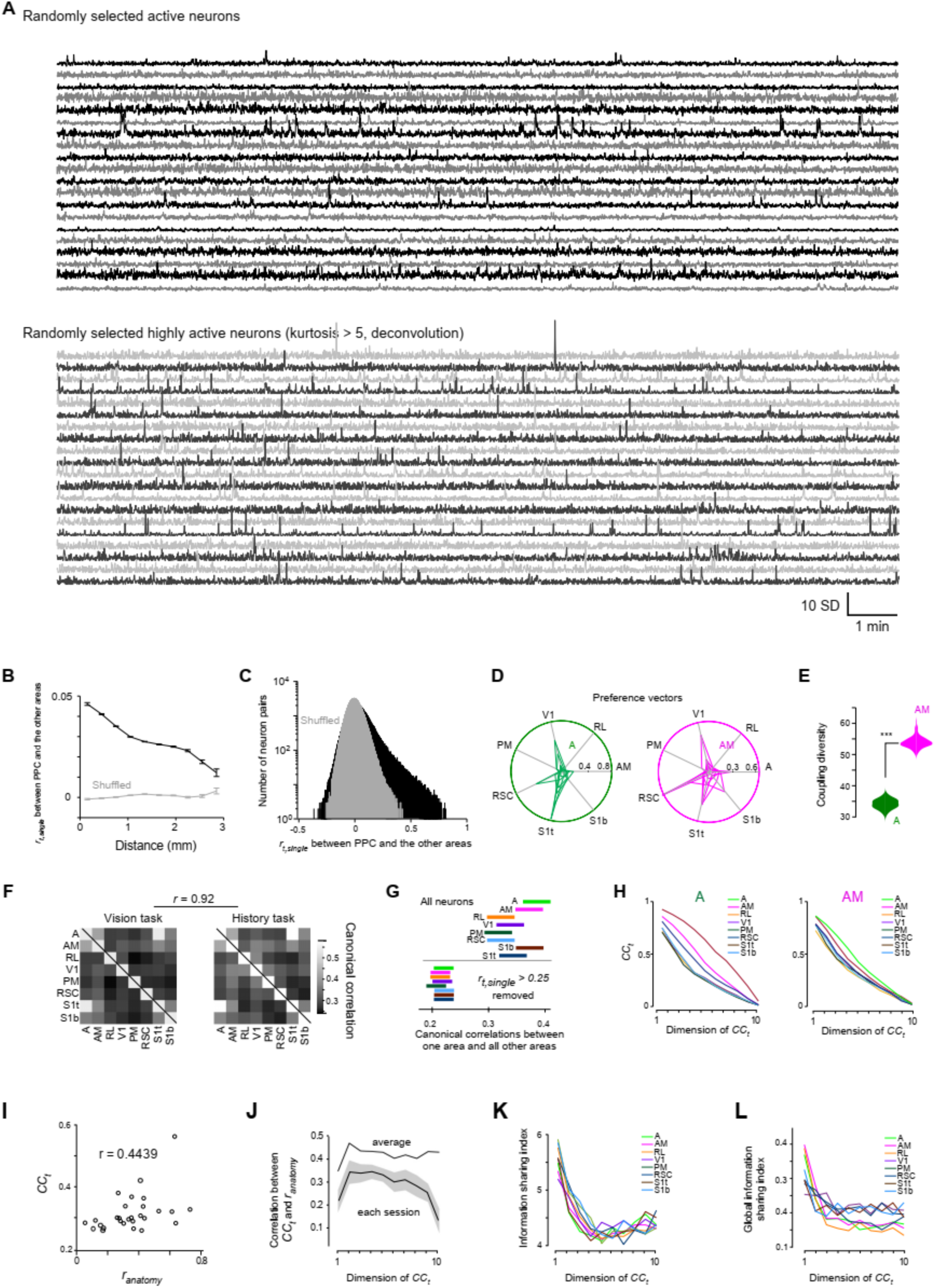
**A.** Randomly selected 20 active neurons (top) and randomly selected 20 highly active neurons (kurtosis > 5, bottom). **B-J.** The same analysis using only highly active neurons (**B: Figure5A.; C: Figure5B; D: Figure5E** (*r_t,single_* >0.25)**; E: Figure5F; F: Figure5H; G: Figure5I; H: Figure5J, I: Figure6L; J: Figure6M, K: Figure6N, L: Figure6O**).

## Supplementary movies.

**Movie S1.**

Two-photon calcium imaging of mouse #1 during task performance. Left, entire field of view (3 mm ×3 mm). Right, square portion of image shown in the right (0.63 mm × 0.63 mm). After registration, a Gaussian filter with standard deviation of 0.36 s was applied to each pixel in the temporal direction for visualization. Time (s) is shown in the upper left. The first frame displays the average image and the location of the cortical areas. The original image was 1345 x 2500 pixels, taken at 5.5 fps. Raw data will be available in https://figshare.com.

**Movie S2.**

Natural images used as visual stimuli for head-fixed mice. A small camera was placed in a mouse cage at the mouse’s eye level to mimic the mouse’s field-of-view. Moving the camera produced global motion and moving the mouse in the cage provided local motion. The luminance of the frame was averaged across the video frames. The video was presented in grayscale at 30 frames/s.

**Movie S3.**

Trial-average image (3mm x 3mm) of mouse #1 for video stimulus + left choice trials (left, magenta) and video stimulus + right choice trials (center, green). The images were aligned at the onset of visual stimulation. Time (s) was shown in the upper left. Visual stimulus onset was set to zero seconds. In the right panel, these two images were overlaid. During visual stimulation, there is an activity sequence of white cells around the visual cortices, which is related neither to left nor right choices, but is the activity that was evoked similarly by the video frames. Images around 4-s included selective activities for left (magenta) and right (green) choices. Note that in the AM, white cells for vision and green and magenta cells for choice are all mixed together. After registration, the trial-average image was obtained, then each pixel was divided by the average intensity of the pixel. Then, the pixel values less than 5 % above the baseline were set to zero for visualization. No filter was used. The first frame shows the average image and the location of the areas. The original image was 1345 × 2500 pixels, taken at 5.5 fps.

